# Epitranscriptomic profiling of VSMC phenotypes reveals uridine modifications linked to post-transcriptional regulation

**DOI:** 10.64898/2025.12.19.694735

**Authors:** Tobias Reinberger, Ayat Ismail, Torben Falk, Janina Fuß, Anja Wiechert, Elke Hammer, Tanja Zeller, Inken Wohlers

**Author notes:** Correspondence to Research Center Borstel, Leibniz Lung Center, Parkallee 1, 23845 Borstel, Germany, +49 4537 / 188-6720.

## Abstract

**Background:** Vascular smooth muscle cells (VSMCs) phenotypic plasticity can modulate atherosclerosis progression. Although several gene regulatory steps towards pro-inflammatory phenotypes have been well-studied, epitranscriptomic changes during this transition and their regulatory roles remain unexplored.

**Methods and Results:** Primary human VSMCs stimulated with TGF-β1 to induce atheroprotective, contractile, and matrix-producing state and with IL1-β plus PDGF-BB to induce a highly energetic, pro-inflammatory state, confirmed by Illumina bulk RNA sequencing and proteomics. Untargeted screening of mRNA base modifications using Oxford Nanopore Technologies direct RNA sequencing and xPore analysis revealed enhanced uridine modification within a GUUUU motif in pro-inflammatory VSMCs. Modified uridines were enriched in 3’-UTR and accessible RNA structures, with implications on Poly(A) tail dynamics and miRNA binding.

**Conclusions:** Atheroprotective and pro-atherogenic treatments induce distinct epitranscriptomic landscapes composed of different modification types, often co-localized in the transcript. Modified uridines in mRNAs are abundant in a high-energy, pro-inflammatory VSMC state and associated with post-transcriptional regulation. n summary, epitranscriptomics adds a novel regulatory layer to VSMC phenotypic transitions critical for atherosclerosis development and progression.

Graphic Abstract

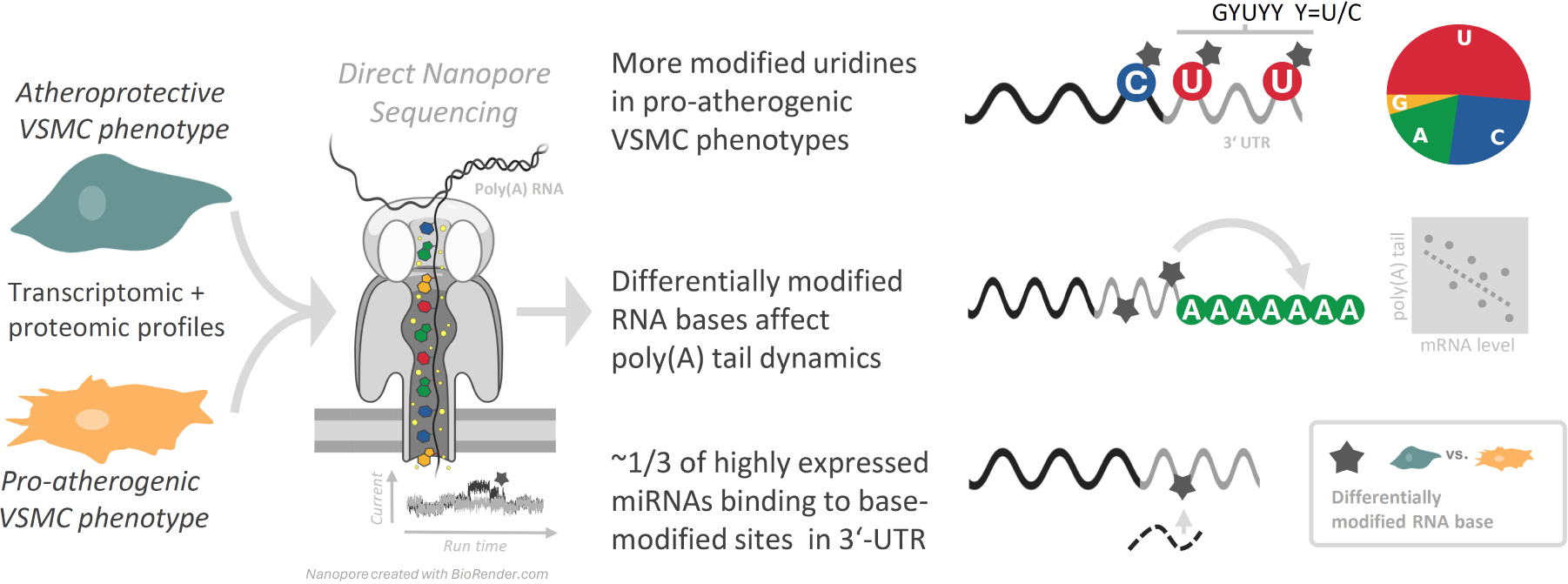

## Introduction

Vascular cell adaptation within atherosclerotic plaques is one of the key features that modulate disease progression and can define the severity of atherosclerotic outcomes.^1^ In particular, the immense plasticity of vascular smooth muscle cells (VSMCs) has been one of the research focuses in cardiovascular disease over the last years. While cytokine-driven pathways, key transcription factors, epigenetic mechanisms, and non-coding RNAs are well described to shape VSMC phenotypic transitions^2–5^ the influence of epitranscriptomic regulation remains largely unexplored and is only beginning to receive attention. ^6^

Epitranscriptomics characterizes and quantifies the collective set of chemica modifications of RNA. Previously, such modifications were investigated with techniques only dedicated to specific modification types, such as N6-methyladenosine (m^6^A) and RNA types such as rRNA or tRNA ^7^ With technological developments in third-generation sequencing, epitranscriptome-wide investigations are becoming feasible. Thereby, direct RNA sequencing of Oxford Nanopore Technologies (ONT) allows the hypothesis-free identification of modification sites by determining significant electric current differences between modified and unmodified bases, measured while RNA passes through a nanopore.^8^

More than 100 different types of RNA modifications have been described, and associations with many diseases have been reported.^9^ Technology-fueled increased interest resulted in a few initial enzyme or modification-centered investigations in vascular disease published within the past five years.^10,11^ The importance of epitranscriptomic regulation in VSMC phenotypic transitions is in the early stages of being recognized. For example, platelet-derived growth factor (PDGF)-BB-induced adenosines to inosines (A-to-I) editing has been described to foster the transition from the contractile, quiescent to a synthetic, pro-inflammatory VSMC state.^12^ Furthermore, it has been demonstrated that ^6^A mRNA methylation stabilizes transcripts of genes nvolved n VSMC-mediated vasoconstriction^13^ and promotes VSMC proliferation, migration, and phenotypic switching^14^ thereby highlighting the clinically relevant role of epitranscriptomic control in blood pressure and neointimal hyperplasia. Another disease-relevant RNA modification is 8-oxo-guanosine (8-oxoGuo), a product of RNA oxidation, which has been associated with atherosclerosis and reported as a predictor of mortality in individuals with type 2 diabetes.^15^ Although the molecular consequences of RNA guanosine oxidation are well described in the context of cardiovascular diseases^16^ the direct implication in VSMC biology remains elusive. Nevertheless, oxidative stress is widely accepted to play a key role in chronic inflammation, to contribute to VSMC phenotypic switching^17^ and to promote the development of unstable plaques^18^

Here, we employed hypothesis-free ONT direct RNA sequencing to extend the spectrum of SMC-relevant RNA modifications beyond ^6^A and to obtain an unbiased view of the global epitranscriptomic landscape across distinct VSMC phenotypic states. Inspired by recent study in which ONT long-read cDNA sequencing has been applied to investigate alternative splicing of contractile and proliferative, pro-inflammatory phenotypes, represented by transforming growth factor β1 (TGF-β1)- and PDGF-BB-treated human aortic smooth muscle cells^19^ we treated VSMCs with TGF-β1 to promote an atheroprotective state (later referred to as condition C3). This VSMC phenotype was compared to pro-inflammatory/pro-atherogenic VSMCs under PDGF-BB / interleukin 1β (IL-1β) treatment^20^ (condition C4), and additionally to oxidative stress-prone VSMCs (C2) as well as oxidatively stressed VSMCs (C1). Here, we demonstrate that epitranscriptomic alterations drive post-transcriptional regulation by affecting mRNA processing, miRNA binding, and translation, thereby determining VSMC states important for atherosclerotic disease progression.

## Methods

Key result files, scripts, and figures have been made publicly available at FAIRDOMHub under “OMICS of vascular smooth muscle cells under oxidative stress” and can be accessed at https://fairdomhub.org/projects/461 Transcriptome sequencing data (Illumina mRNA and small RNA-Seq as well as ONT direct RNA-Seq) for this study have been deposited in the European Nucleotide Archive (ENA) at EMBL-EBI under study accession number PRJEB98980 (https://www.ebi.ac.uk/ena/browser/view/PRJEB98980). Proteomics data files are available in PRIDE under accession ID PXD065513, currently accessible via reviewer access token w5MHzwMvIHve. The modification workflow is available at https://github.com/iwohlers/2025_ont_rnaseq_vsmc A detailed method description, Supplementary Figures, Supplementary Tables, and Major Resource Tables are available in the Supplemental Material.

### Cell culture and functional assays of vascular smooth muscle cells

Human primary vascular smooth muscle cells (VSMCs) were obtained from multiple commercial vendors (Thermo Fisher Scientific/Gibco, Cell Applications, Inc., and PromoCell), all of whom adhere to the principles of the Declaration of Helsinki (see **Supplementary Table S1**). VSMCs were cultured and expanded up to passage eight. Since VSMCs rapidly transform to a synthetic phenotype if seeded on a plastic surface, we cultured VSMCs on a collagen hydrogel instead. In brief, cells were seeded onto collagen hydrogels and allowed to attach for 24 h in medium containing 1% fetal bovine serum (FBS) and µM cytochalasin D to inhibit contraction, which critical to VSMC monolayer for CellROX measurements. A contractile, atheroprotective phenotype was induced by treatment with 5 ng/mL TGF-β1 and µM cytochalasin D for four days, followed by induction of a pro-inflammatory/ pro-atherogenic phenotype using 10 ng/mL PDGF-BB and 10 ng/mL IL-1β for two days. Cells were subsequently used for CellROX assays and RNA sequencing. Detailed culture conditions and hydrogel preparation are provided in the Supplementary Material. A detailed description of functional assays such as oxidative stress assays, quantification of the mitochondrial membrane potential and seahorse assays are also provided in the Supplementary Material.

### Short-read RNA sequencing

Short-read sequencing using the TrueSeq stranded mRNA library preparation kit performed for eight biological replicates (see **Suppl. Table S1**) and five conditions oxidatively-stressed VSMCs (C1; TGF-β1 and PDGF-BB / IL-1β PDGF boost), oxidative stress-prone VSMCs (C2; TGF-β1 and PDGF-BB / IL-1β), atheroprotective VSMCs (C3; TGF-β1 only), proatherogenic VSMCs (C4; PDGF-BB / IL-1β only), and untreated VSMCs (C5). RNA isolation and library preparation is descirped in the Supplementary Material. RNA sequencing (RNA-Seq) datasets were analyzed using a community workflow from the snakemake workflow catalog (https://github.com/snakemake-workflows/rna-seq-kallisto-sleuth) version 2.8.4. A detailed descrition of the workflow tools, which cover the enrichment analysis and isoform switch analysis is provided in the in the Supplementary Material. Principal component analysis (PCA) performed for the 1,000 most variable genes after selecting the most highly expressed transcript per gene and keeping only transcripts with a minimum normalized log_2_ expression of 6.

### Long-read direct RNA sequencing by Oxford Nanopore Technologies

Stimulated VSMCs cultured on Col hydrogels were subjected to direct RNA-Seq using Oxford Nanopore Technologies (ONT) (**Suppl. Table S1**) und four the conditions C1 C4 (see above). To trigger endogenous ROS production, VSMCs were incubated with 400 ng/mL PDGF-BB for 3 hours. RNA isolation was performed as described above for short-read RNA-Seq. Approximately 10×10^6^ stimulated cells per condition were used as input for direct RNA-Seq. Library preparation and direct RNA-Seq were conducted according to the ONT protocol SQK-RNA002. About 150–200 ng rRNA-depleted RNA was sequenced for 18 22 hours uisng a GridION device with FLO-MIN106 flow cells with more than 1000 active pores. ONT direct RNA-Seq data was processed using our own snakemake workflow (in detailed described in the Supplementary Material and available at https://github.com/iwohlers/2025_ont_rnaseq_vsmc), with tools installed in reproducible fashion via conda using environment files provided as part of the workflow. The workflow uses guppy version 6.3.2 for basecalling and pycoQC^21^ for quality control. Transcript expression levels were determined using nanocount^21^ (transcript annotation from Ensembl v113). For differential modification detection, reads were mapped to transcript reference sequences (Ensembl v113) using minimap2^22^ followed by xPore^8^ (version 2.1) used with nanopolish eventalign output, *i.e.* modification positions refer to transcript coordinates. Poly(A) tail lengths were estimated using nanopolish mode ‘polya’ (coverage>10). Coordinates of untranlasted regions (UTRs) and coding sequences (CDS) were obtained by pybiomart v0.2.0. Normalization of nanocount counts and testing of differential expression were performed with DESeq2^23^ and a additional variance stabilization using rlog transformation performed before principal component analysis, both previously described for nanocount expression data.^2^ These downstream analyses and visualizations are available as part of the FAIRDOMHub under https://fairdomhub.org/projects/461

### Transcriptome-wide prediction of RNA structure elements

RNA secondary structure prediction and analysis were performed in Python using ViennaRNA and Forgi packages. For each modified base or k-mer, overlapping 150-nt mRNA sequence windows were extracted from Ensembl annotations and shifted stepwise across the modification site Centroid secondary structures and minimum free energies (MFE) were predicted for each window. Structural similarity was quantified from dot–bracket representations using a sequence alignment–based scoring approach and used to cluster centroid structures by hierarchical clustering. Representative structures were selected based on cluster size and energy characteristics, and the lowest-MFE centroid structures were retained for downstream analysis. Secondary structure elements were annotated using Forgi, and enrichment of structural elements within k-mers was assessed using hypergeometric and Fisher’s exact tests (for details see Supplementary Material).

### Prediction of RNA structure elements with miRNA binding

Wide-window prediction (>350 nt) of RNA secondary structures was conducted with RNAplfold (version 2.7.0). Alignment of the miRNA involved two steps: local alignment of the seed sequence and global upstream alignment of the residual miRNA sequence (for details see Supplementary Material).

### Small RNA sequencing

For the identification of highly expressed miRNAs, small RNA sequencing was performed for two technical replicates of four conditions (C2 C5), respectively. Approximately 3.8×10^5^ cells/well (HAOSMC IV) seeded in 6-well plates that covered with Col hydrogel (1.5 mg/mL). VSMCs were stimulated as described above for short-read RNA-Seq and RNA isolation, library preparation, and data analysis are described in the Supplementary Material. We investigated whether the modified base is part of miRNA seed regions, for which we considered 7mer-m8 sites, *i.e.* bases 2-8 of the miRNA need to match the complement of the region covering the modified base Seed regions obtained from Targetscan^24^ Annotation of publication-supported miRNA-target interactions was obtained from MirTarBase^25^

### Proteomics

Protein quantification in TGF-β1– and PDGF-BB–stimulated VSMCs was performed by label-free quantitative mass spectrometry (MS) with four technical replicates per condition. Proteins were extracted, quantified, enzymatically digested, and analyzed by HPLC–ESI–MS/MS using an Orbitrap mass spectrometer. MS data were processed and normalized with Spectronaut using a human UniProt reference database, and protein abundances were calculated as maxLFQ values. Statistical analysis was performed at the peptide level and protein set enrichment analysis conducted using Enrichr^26^ Detailed sample preparation, acquisition parameters, and analysis workflows are provided in the Supplementary Material.

## Results

### Energetic and transcriptomic profiling of VSMC phenotypes

To study pro-atherogenic transitions of vascular smooth muscle cells with respect to oxidative stress, inflammatory signaling, gene regulation, and epitranscriptomic changes, we established an *in vitro* cell culture model and performed energetic and global transcriptomic profiling of these VSMC phenotypes (**Figure 1A Suppl. Figure S1A,** and **Suppl. Table S2**). As previously described^27^ activation of TGF-β signaling promoted a quiescent, contractile-like VSMC phenotype, accompanied by reduced mitochondrial activity (**Suppl. Figure S1**), higher level of canonical SMC contractile marker genes (*e.g. ACTA2 CNN1 TAGLN* and *LMOD1 cf.* **Suppl. Table S3**), and activation of genes involved in extracellular matrix organization (ECM) (**Figure 1**). This mesenchymal-like, ECM-producing VSMC phenotype can contribute to fibrous cap formation in atherosclerotic plaques^3^ protective layer that stabilizes fatty deposits in the vessel wall and helps prevent plaque rupture. Therefore this VSMC phenotype s referred to as the atheroprotective VSMC state or C3 throughout the study.

**Figure 1.**
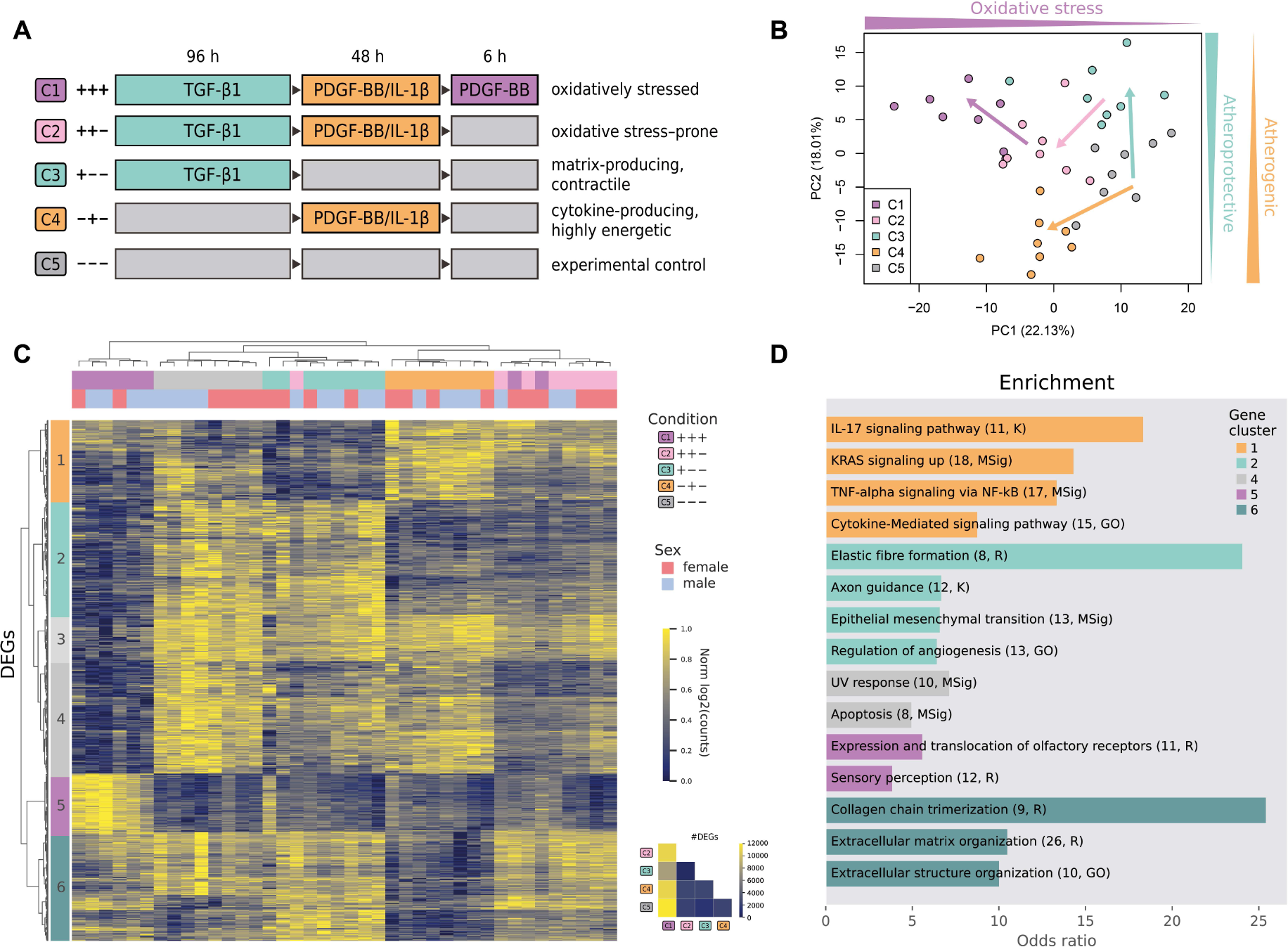
(**A**) Experimental design and definition of conditions C1–C5 used throughout the study. Each condition is sampled with eight biological replicates. (**B**) Principal component (PC) analysis of five conditions is based on the 1,000 most variable transcripts with normalized and batch-corrected log_2_ expression ≥ 5. (**C**) Clustered heatmap showing row-scaled expression of the 1,000 most variable canonical, differentially expressed transcripts (with FDR<=0.001; mean log_2_ count ≥ 5) from the joint model across all conditions. (**D**) Up to four top enriched pathways for six gene clusters displaying condition-specific up- or down-regulation, colored according to the corresponding condition.

PCA of the most variable expressed genes showed the largest variation (PC1) between atheroprotective or untreated VSMCs (C3/C5) and PDGF-BB / IL-1β–treated VSMCs (C1/C2/C4) (**Figure 1B, Suppl. Figure S2**). Of note, we added IL-1β as it can synergistically promote PDGF signaling.^20^ PDGF-BB / IL-1β–treated VSMCs (C4) exhibited an energetic and pro-inflammatory signature, with enhanced mitochondrial and glycolytic activity as well as increased expression of genes nvolved n cytokine-induced signaling (**Suppl. Figure S1 and Figure**). In particular IL-17, TNF/NF-κB, and RAS signaling pathways were shown to worsen atherosclerosis and accelerate plaque progression.^28,29^ Accordingly, PDGF-BB / IL-1β–treated VSMCs (C4) are hereafter termed pro-atherogenic VSMCs.

In addition, we created an intermediate VSMC state (TGF-β1 PDGF-BB / IL-1β; C2) between atheroprotective and pro-atherogenic phenotypes, with mesenchymal transition–associated genes remaining inactive (cluster 2) while extracellular matrix and inflammatory genes are upregulated (clusters and 6 in **Figure 1C**). Notably, impairment of mesenchymal transition programs is often associated with reduced cellular adaptability to stress.^30^ Moreover, the concurrent production of ECM components and cytokines is energy-demanding and is therefore associated with increased ROS levels, which can promote mitochondrial dysfunction and overwhelm cellular antioxidant defenses. The intermediate state is therefore termed oxidative stress–prone VSMCs.

Finally, we added very high levels of PDGF-BB to this oxidative stress–prone VMSC phenotype characterized by elevated biosynthetic activity and reduced stress adaptation capacity, since we suspected high local concentrations of PDGF in atherosclerotic lesions ^31^ Indeed, hyperactivation of PDGF signaling resulted not only in elevated ROS levels (**Suppl. Figure S1B**), most likely through activation of mitochondria and NADPH oxidase as previously described^32^ but also led to reduced expression of several genes essential for cell function, accompanied by strong upregulation of olfactory receptor genes (**Figure 1D**), which were found to regulate pathophysiological processes in SMCs and described as novel therapeutic targets.^33^

In addition to pronounced transcriptional changes (**Supplementary Fig. S3**), oxidative stress induced widespread differential splicing. A total of 1,298 genes exhibited significant splicing differences between atheroprotective VSMCs and pro-inflammatory phenotypes, with CCL20, CXCL8, FAH, and NOL4L consistently affected across all comparisons. For example, increased CCL20 expression in pro-atherogenic VSMCs was accompanied by preferential usage of isoforms containing the chemokine interleukin-8-like domain (**Suppl. Figure S4A**).

### Epitranscriptomic profiling of VSMC phenotypes

In order to investigate whether pro-inflammatory conditions accompanied by high level of ROS can directly modulate mRNA base modifications, we used Oxford Nanopore (ONT) direct RNA sequencing and xPore to obtain a comprehensive overview of the RNA modification landscape. In general, up to 4% of mRNA bases of an individual transcript were found to be modified (*e.g.* methylated; rate ≥ 0.5) in atheroprotective VSMCs (**Figure 2A** and **Suppl. Table S5**). While the absolute number of modified bases per transcript increases with transcript length, the ratio of modified bases per transcript correlates positively with ONT mRNA abundance but negatively with transcript length. Thus, shorter mRNAs tend to have a higher density of modified bases while expressed at higher levels.

**Figure 2:**
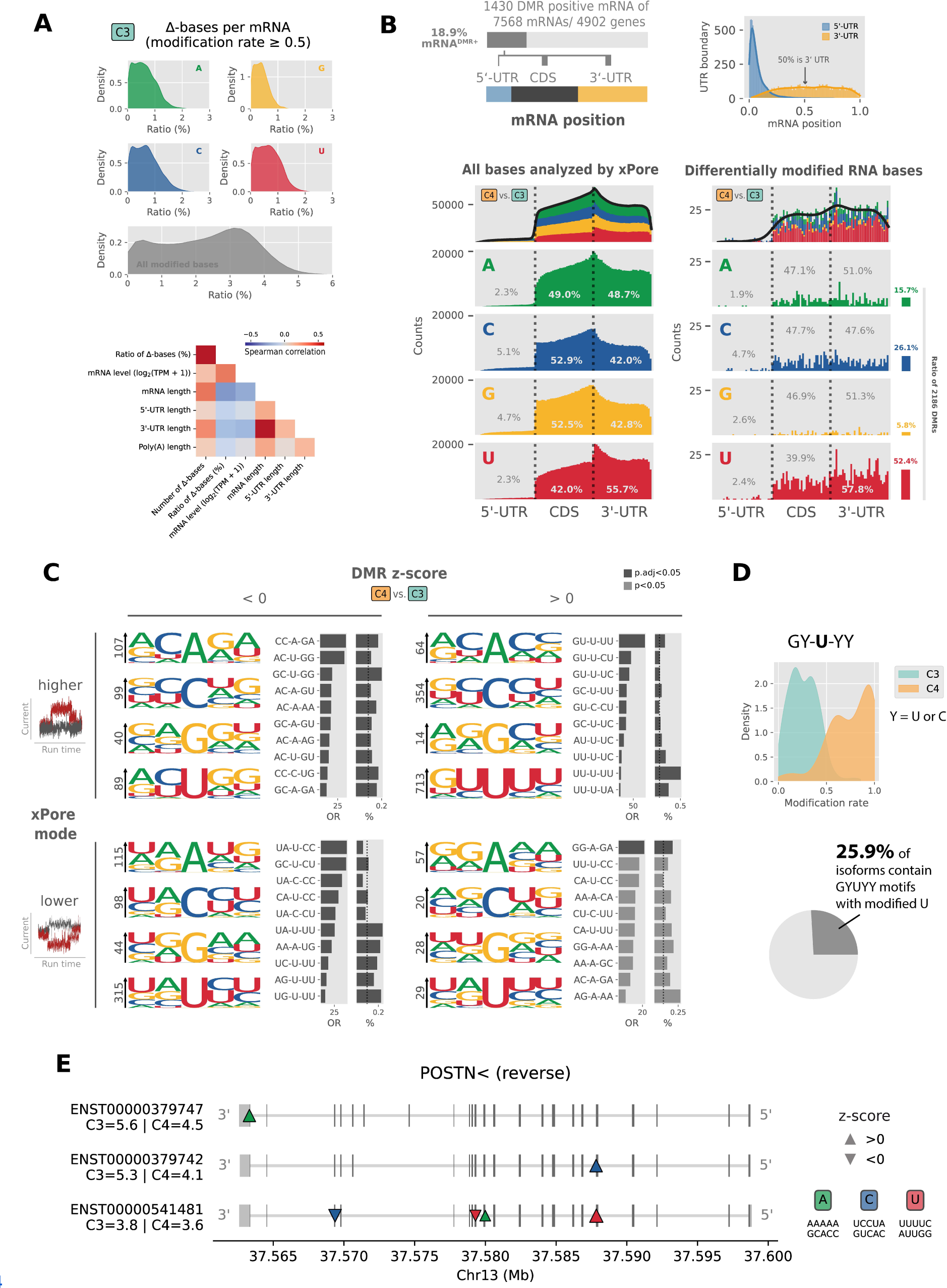
Detection of RNA base modifications by ONT direct RNA sequencing and xPore in vascular smooth muscle cells. VSMCs were treated with atheroprotective TGF-β1 (C3) or pro-inflammatory PDGF-BB / IL-1β (C4) (n=3 technical replicates). Approximately 4 million base positions across 7,568 transcripts were analyzed. (**A**) The proportion of modified (Δ) RNA bases (rate ≥ 0.5) in TGF-β1–treated VSMCs peaks at 2 to 4%. A Spearman correlation matrix is shown below. (**B**) A total of 2,186 significantly differentially modified RNA bases (DMRs) were identified in UTRs and coding sequences (CDS) across 1,430 transcripts. The relative distribution of RNA bases (A, C, G, U) within UTRs and CDS is shown for all analyzed positions (left) and for significant DMRs (C4 vs C3; right). (**C**) Sequence motif analysis highlights the GUUUU motif with higher modification rates of uridine in C4 (DMR z-score > 0). The xPore mode “higher” indicates increased Nanopore current at modified bases The odds ratio (OR) and percentage (%) of the 10 most abundant sequence motifs (5-mers) are shown next to the motif plots. (**D**) Distribution of modification rates for the central uridine in GY–U–YY motifs (adjusted *p* < 0.05; rate ≥ 0.25), representing ∼26% of all DMR-positive mRNAs. (**E**) *POSTN* transcripts illustrate isoform-specific DMRs.

Unexpectedly, we did not find evidence that the endogenous induction of oxidative stress is associated with increased RNA oxidation represented by 8-oxoGuo. Moreover, we found the lowest number of modified mRNA sites (n=1,220) when comparing atheroprotective VSMCs (C3) with oxidatively stressed VSMCs (C1) (**Suppl. Table S8**).

However, comparison of atheroprotective VSMCs (C3) and pro-atherogenic VSMCs (C4) yielded the largest number of detectable mRNA modification sites (2.1 million in C3 vs C4), from which 5,989 high-confidence differentially modified RNA bases (DMRs) were identified after multiple-testing correction (FDR ≤ 0.05; **Suppl. Table S6** and **S7**). Subsequent filtering for a differential modification rate of at least 25% kept 2,186 differential sites. These DMRs most likely reflect multiple modification types affecting adenosine, cytidine, and uridine (**Suppl. Figure S5**) which appear to be cytokine-dependent rather than stress-induced.

We observed that PDGF-BB treatment led predominantely to enhanced modification of uridines in pro-athorogenic VSMCs (PDGF-BB / IL-1β; C4) as well as oxidatively stressed VSMCs (TGF-β1 PDGF-BB / IL-1β PDGF-BB boost; C1) (**Figure 2B-D** and **Suppl. Figure S6**). The highest number of modified uridines were found in highly energetic, metabolic active and cytokine-produncing VSMCs. Uridine modifications often occurred in GY-U-YY motifs, with Y being a pyrimidine base (C or U). Thereby, GUUUU was the most prevalent 5-mer context (209 DMR sites in 194 transcripts), showing ∼80-fold enrichment across all mRNA sequences analyzed with xPore (**Figure 2C**). Affected genes appear to be regulated by the transcription factors TCF-21, JUN (c-Jun) or JunD and are enriched for epithelial-mesenchymal transition, immune system, cell migration, and the PI3K/AKT/mTOR signaling pathway (**Suppl. Table S9**). Analysis with xPore revealed that many transcripts harbor multiple differentially modified sites, and co-modification of uridine together with adenosine and/or cytidine was frequently observed (cf. **Suppl. Table S10**). Overall, the comparison of atheroprotective VSMCs (C3) and pro-atherogenic VSMCs (C4) yielded 1,258 genes expressing transcripts that contain one or more DMR-sites, with the top affected genes *FN1* (18 sites, 9 transcripts), *ITGB1* (15 sites, 12 transcripts), *HDLBP* (13 sites, 3 transcripts), and *TUBB* (10 sites, 5 transcripts) (cf. **Suppl. Table S11**). Of note, *FN1* and *ITGB1* are described to be major regulators of VSMCs.^34^ Other DMR-affected genes and transcripts involved in VSMC-relevant processes are highlighted in **Suppl. Table S12** and **Suppl. Figure S7** Except *COL6A2* and *PXDC1* (**Suppl. Figure S4B** and **S4C**), of these notably DMR-affected genes is among the 272 genes for which significant differential splicing was detected between atheroprotective and pro-atherogenic VSMCs (**Suppl. Table S4**), suggesting that DMRs may not affect alternative splicing.

### Association of base modification with protein abundance

Since we detected the largest number of DMRs when comparing atheroprotective (TGF-1β; C3) with pro-atherogenic VMCs (PDGF-BB / IL1-β1; C4), we subsequently quantified the proteome of these VSMCs phenotypes (**Suppl. Table S13**), resulting in 4,995 dentified proteins of which 1,171 were significantly differentially abundant after multiple testing correction (**Suppl. Table S14**). Enrichment analysis confirms the pro-inflammatory signature of PDGF-BB / IL1-β1-treated VSMCs (**Figure 3A**). In order to investigate a potential role of RNA modifications on protein levels, we focused on transcripts detected at a minimum of 10 transcripts per million (TPM) in both Illumina and ONT sequencing. At this threshold, log10-transformed RNA levels derived from Illumina and ONT correlate best, with a Pearson’s correlation coefficient of r=0.66 for C3 and r=0.67 for C4 (cf. **Suppl. Figure S8**), which is within ranges reported previously^35^ In addition, transcript counts were aggregated per gene since proteomics is not isoform-sensitive. Protein levels align with gene expression levels in atheroprotective VSMCs (TGF-1β; C3) when only non-zero values are considered, and this relationship is independent of the presence of differentially modified RNA bases (DMRs in C4 vs C3). However, DMRs are more frequently identified at higher gene/protein levels (**Figure 3B**). Notably, the proportion of DMR-positive mRNAs decreases from approximately 22% among mRNAs with detectable protein to about 8% among those for which no corresponding protein was detected despite comparable mRNA levels suggesting that some DMRs may be crucial for translation activity. Moreover, for genes linked to more than three DMRs, changes in transcript abundance were strongly mirrored at the protein level. This is reflected by strong correlation between the log fold change in gene expression and the corresponding log_2_ fold change in protein abundance. (**Figure 3C**). This relationship supports a role for DMRs in shaping transcriptional programs that extend to protein expression, particularly for genes linked to the TCF21-regulated atheroprotective, fibroblast-/mesenchymal-like VSMC state (C3). A subset of these genes is downregulated during the transition to the pro-inflammatory VSMC phenotype (C4). An overview of genes with notable differences in corresponding protein levels is shown in **Figure 3D** Neither the relative position of DMRs within UTRs or coding sequences nor the direction of modification changes suggests a consistent rule governing DMR-dependent transcriptional or translationa regulation. Nonetheless, we observe a trend toward reduced adenosine modification (z-score 0) in genes exhibiting higher protein abundance in pro-atherogenic VMCs, and the opposite trend in genes with lower protein abundance. Moreover, cytidines appear to be differentially modified more frequently when located in the 5’-UTR of genes associated with increased protein levels.

**Figure 3.**
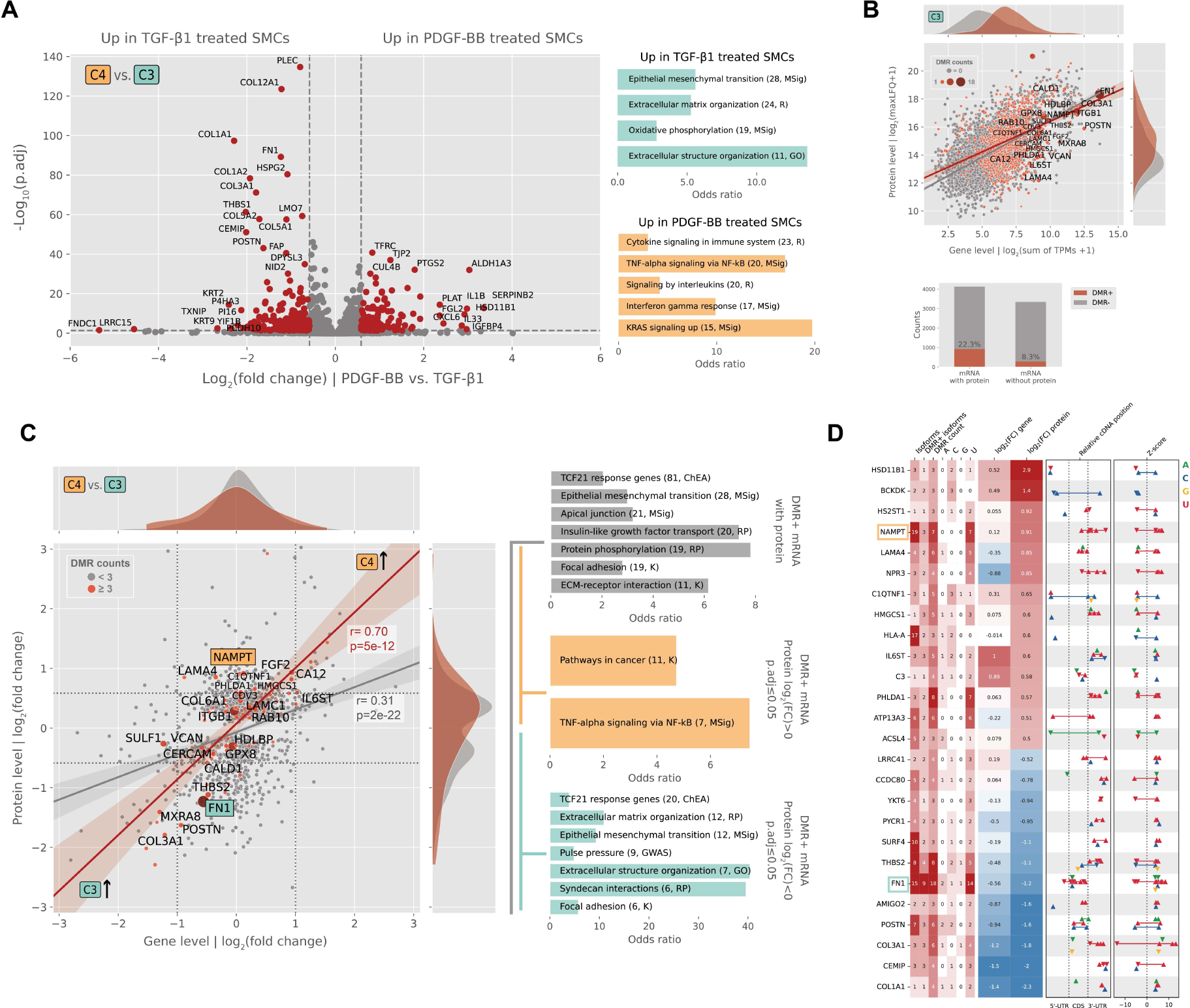
Impact of RNA base modifications on translational activity in smooth muscle cells. (**A**) Differential proteomic analysis of VSMCs treated with TGF-β1 (C3) or PDGF-BB / IL-1β (C4) (n = 4 technical replicates per condition). Volcano plot and enrichment analyses highlight canonical TGF-β1– and PDGF-associated proteins and pathways (GO=Gene Ontology; MSig=Molecular Signature database; R=Reactome database). (**B**) Regression analysis of gene-level versus protein-level expression in C3 for transcripts with (red) or without (gray) differentially modified RNA bases (DMRs; C3 vs C4). Point size reflects the number of DMRs per gene across all isoforms. (**C**) Same analysis as (**B**) using log_2_ fold changes (C3 vs. C4). The Pearson correlation (r; significance assessed with t-test) was calculated for genes with ≥ 3 DMRs across all isoforms and < 3 DMRs. Gene set enrichment analysis was performed for genes with ≥ 3 DMRs and stratified by corresponding protein log fold change (> 0 or < 0). (ChEA=ChIP-X enrichment analysis; GWAS= GWAS catalog; K; KEEG database; RP Reactome Pathways). (**D**) Overview of genes with |log_2_FC| 0.5 (FC=fold change), discordant gene–protein log FC (> 0.25), and > 2 DMRs across all isoforms; including isoform counts, DMR position in untranslated regions (UTR) or coding sequence (CDS), and z-scores (C4 vs C3) with xPore mode as arrows (“higher”=up, “lower”=down).

### Transcriptomic landscape of mRNA modification–related genes

Since uridine was the most frequently modified RNA base, we first examined the expression level of pseudouridine synthase genes in atheroprotective and pro-atherogenic VSMCs. We observed modest yet significant downregulation of three of them, but only in PDFG-treated, oxidatively stressed VSMCs (C1) (**Suppl. Figure S9**). In contrast, a notable increase in *PUS7* level was observed in pro-atherogenic VMCs, reaching nominal significance (p 0.022, p.adj = 0.098) (**Figure 4A**). Among all mRNA modification–related genes (median log_2_ normalized counts > 5; cf. **Suppl. Table S15**), significant differences between atheroprotective and pro-atherogenic VSMCs were detected only for *TET1* (hm5C oxidase) after adjustment for multiple testing (p.adj. 0.05). Although it appears that only a few mRNA modifiers and genes encoding for mRNA-binding proteins differ individually, the global expression signatures clearly separate the two VSMC phenotypes (**Suppl. Figure 4B**), suggesting an orchestrated program of mRNA modification–related genes that promote either atheroprotective or pro-atherogenic pathways.

**Figure 4:**
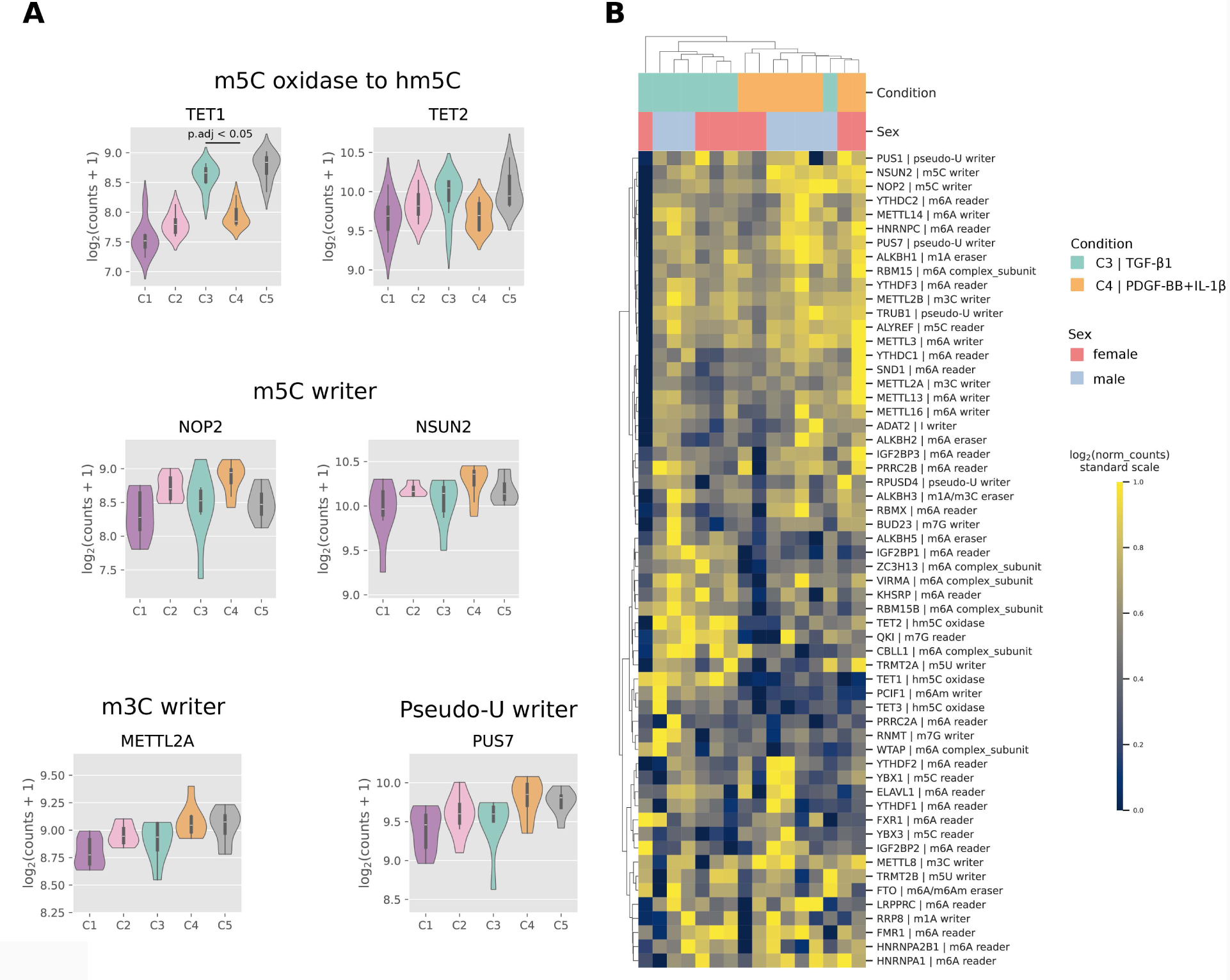
Expression pattern of genes encoding mRNA-modifying enzymes in atheroprotective (C3) and pro-atherogenic (C4) VSMCs. (**A**) Violin plots of genes whose canonical transcripts are differentially expressed in C3 vs C4 at nominal significance (p < 0.05; for TET1: p.adj < 0.05; **Suppl. Table S3**). (**B**) The clustermap shows the standardized expression level of genes with median log2(normalized counts) ≥ 5 (**Suppl. Table S15**).

### RNA base modifications influence poly(A) tail dynamics

Next, we examined how DMRs influence mRNA processing and translationa regulation, including poly(A) tail dynamics. Density plots of poly(A) tail lengths in atheroprotective (TGF-1β; C3) and pro-atherogenic VMCs (PDGF-BB / IL1-β1; C4) reveal that genes/transcripts with increasing numbers of DMRs exhibit progressively longer poly(A) tails, along with more pronounced differences between the two VSMC phenotypes (**Figure 5A**). In addition, we observed a strong correlation between gene/protein log fold changes and corresponding shifts in poly(A) tail length. Longer poly(A) tails appear to be associated with lower mRNA levels as well as protein levels in pro-atherogenic VSMCs (**Figure 5B**). This relationship is highlighted for *NAMPT* and *FN1* in **Figure 5C**, which were the only ones with significant differences in poly(A) tail length after adjusting for multiple testing (cf. **Suppl. Table S16**). While *NAMPT* shows higher expression in pro-atherogenic VSMCs (C4), accompanied by shorter poly(A) tails, *FN1* exhibits the opposite pattern when all isoforms are considered. Interestingly, several uridines show increased modification rates (in “higher” xPore mode) in pro-atherogenic VSMCs. These modified uridines are located predominantly in the 3′-UTRs of *NAMPT* isoforms, whereas in *FN1* isoforms they occur exclusively within coding sequences.

**Figure 5.**
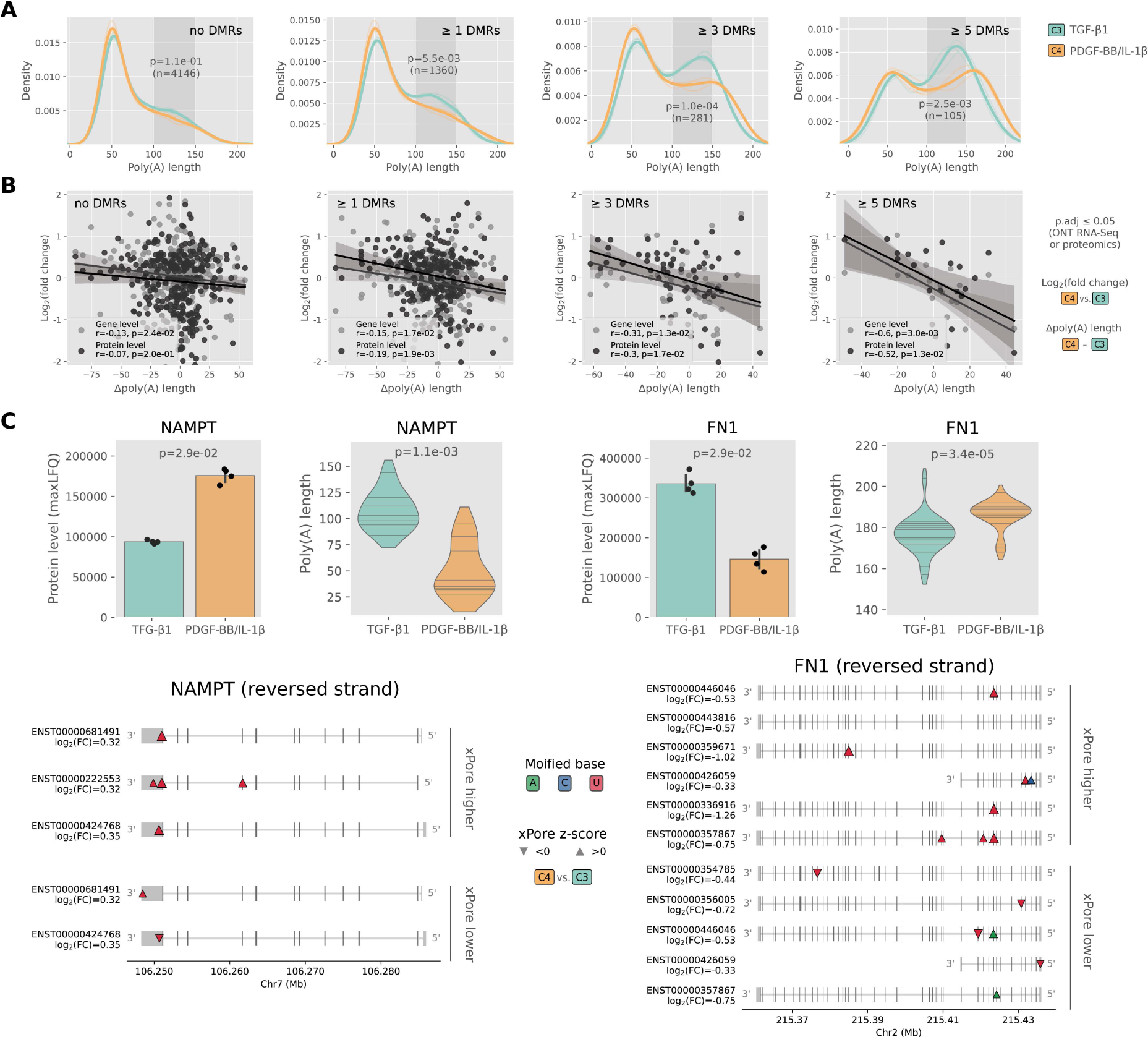
Poly(A) tail length distribution of mRNAs isolated from TGF-β1– (C3) or PDGF-BB / IL-1β–stimulated vascular smooth muscle cells (C4). (**A**) Kernel density plots show increasing poly(A) tail length and greater differences between C3 and C4 with increasing number of differentially modified RNA bases (DMRs). Statistica significance wa calculated using a permutation test in the region 100 to 150 adenosines, with transcripts counts in parentheses. (**B**) Pearson correlation plots of difference (Δ) in poly(A) tail length versu log_2_ fold change for differentially expressed genes/proteins (p.adj ≤ 0.5). Significance assessed with t-test. (**C**) Examples illustrating the relationship between protein abundance (n=4 technical replicates), poly(A) tail length, and DMR position (n=3 technical replicates). Poly(A) tail lengths across all isoforms (polya coverage > 10; FN1 = 9 isoforms, NAPT = 3) and technical replicates (n=3) are shown per gene significance was assessed by Mann–Whitney U tests. Lower panels display isoforms with DMRs stratified by xPore mode (“higher” or “lower”), with log_2_ fold change (C4 vs C3) shown for each transcript.

### Modified uridine bases are enriched in accessible RNA secondary structures in the 3’-UTR

Next, we predicted RNA secondary structures of 150-nucleotide sequences containing DMR sites using the ViennaRNA library to investigate whether DMRs might be found more frequently in distinct structural elements. In brief, 65 controid RNA secondary structures were predicted per transcript and DMR position by shifting a 150 nucleotide window two bases downstream, starting from 140 nucleotides upstream of the DMR position. Centroid RNA secondary structures were clustered based on similarity scores The RNA secondary structure with the lowest MFE of the most representative cluster (selected upon cluster size, median MFE, and MFE variance) was taken for over-representation analysis (**Figure 6A**). As part of our quality control, we examined the negative minimal free energy (MFE) of all predicted centroid RNA secondary structures across relative cDNA positions. Consistent with previous observations, the negative median MFE per position declines toward the 3′-UTR, accompanied by reduced GC content, a hallmark of functional mRNAs^36^ (**Figure 6B**). Over-representation analysis of DMRs n RNA secondary structure elements revealed that differentially modified uridines are 1.7-fold enriched in interior loops in 3’-UTRs (**Figure 6C** and **Suppl. Table S17**).

**Figure 6.**
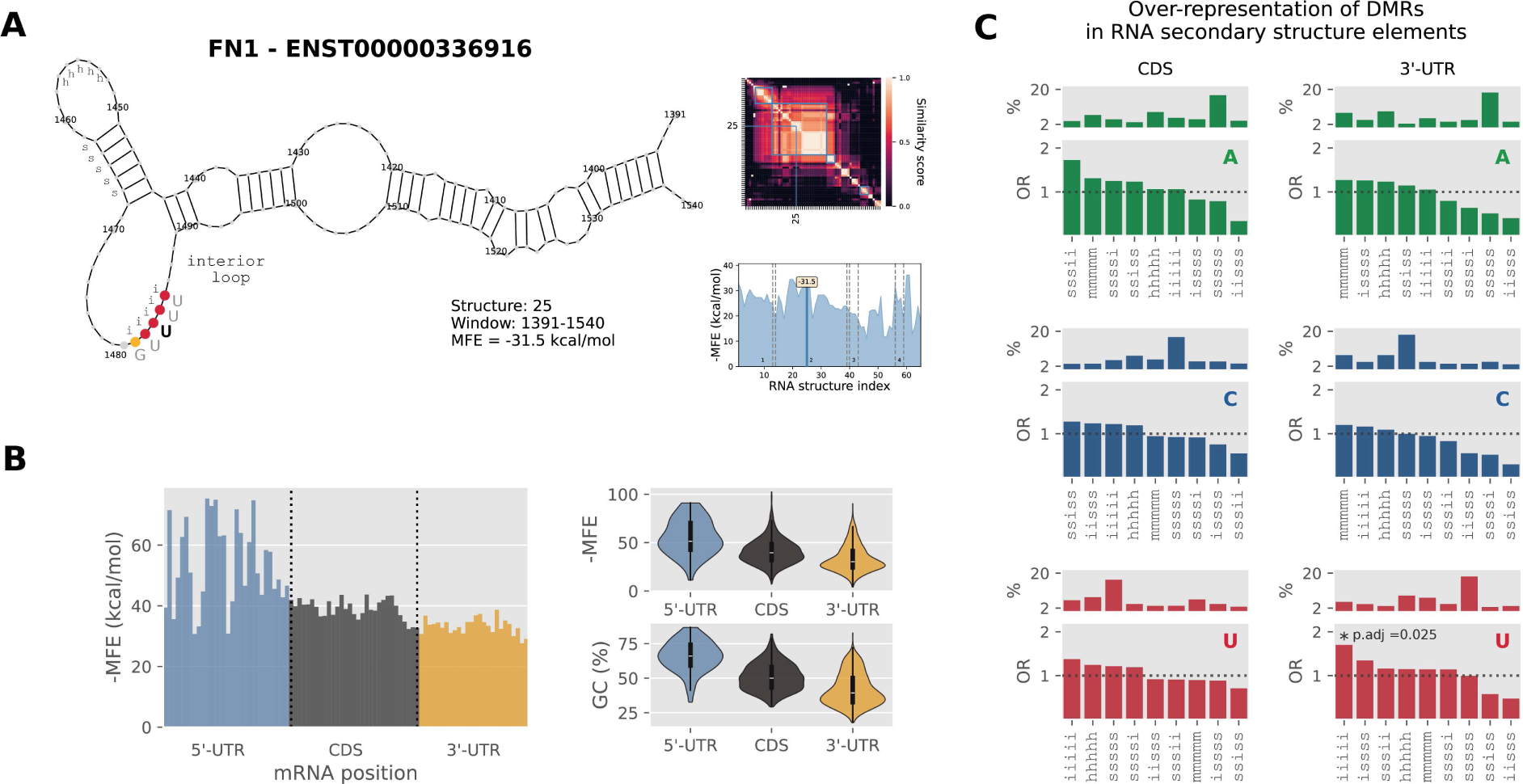
Distribution of differentially modified RNA bases in secondary RNA structures. (**A**) RNA secondary structures surrounding DMRs were predicted from sliding 150-nt windows using the ViennaRNA package. Centroid structures with minima free energy (MFE) were clustered, and dominant secondary structure elements (5-mers) were extracted. (**B**) Mean MFE per binned relative DMR positions (90 bins) in 5’/3’-UTR (untranslated region) and coding sequence (CDS). Violin plots highlight the correlation of MFE and GC-content. (**C**) Differentially modified uridines are enriched in predicted interior loops (iiiii) within 3’-UTR (Fisher’s exact test). Bar plots show relative abundance (%) and odds ratios (OR) of secondary structure elements across 343 A, 571 C and 1,146 U DMRs (*cf.* **Suppl. Table S17**).

### Modified RNA bases may modulate miRNA binding

Moreover, we detected an excess of modifications near stop codons at the beginning of the 3’-UTR (*cf.* **Figure 2B**), which has also been reported previously by the authors of xPore^8^ We hypothesize that this enrichment is related to DMRs that influence miRNA binding, as miRNA-to-mRNA binding is reported to be most effective when occurring at the beginning or the end of long 3′-UTRs.^37^ Therefore, we assessed the potential impact of DMRs on miRNA binding. We quantified miRNA expression in four VSMC states (C2 to C5) and screened highly expressed miRNAs (log_2_(count) ≥ 10 in one replicate) for miRNA-binding sites based on direct 8-mer seed matches We identified 60 of the 202 highly expressed miRNAs whose seed sequences could directly bind to DMR-containing mRNA regions in 3’-UTR (**Figure 7A** and **Suppl. Table S18**).

**Figure 7.**
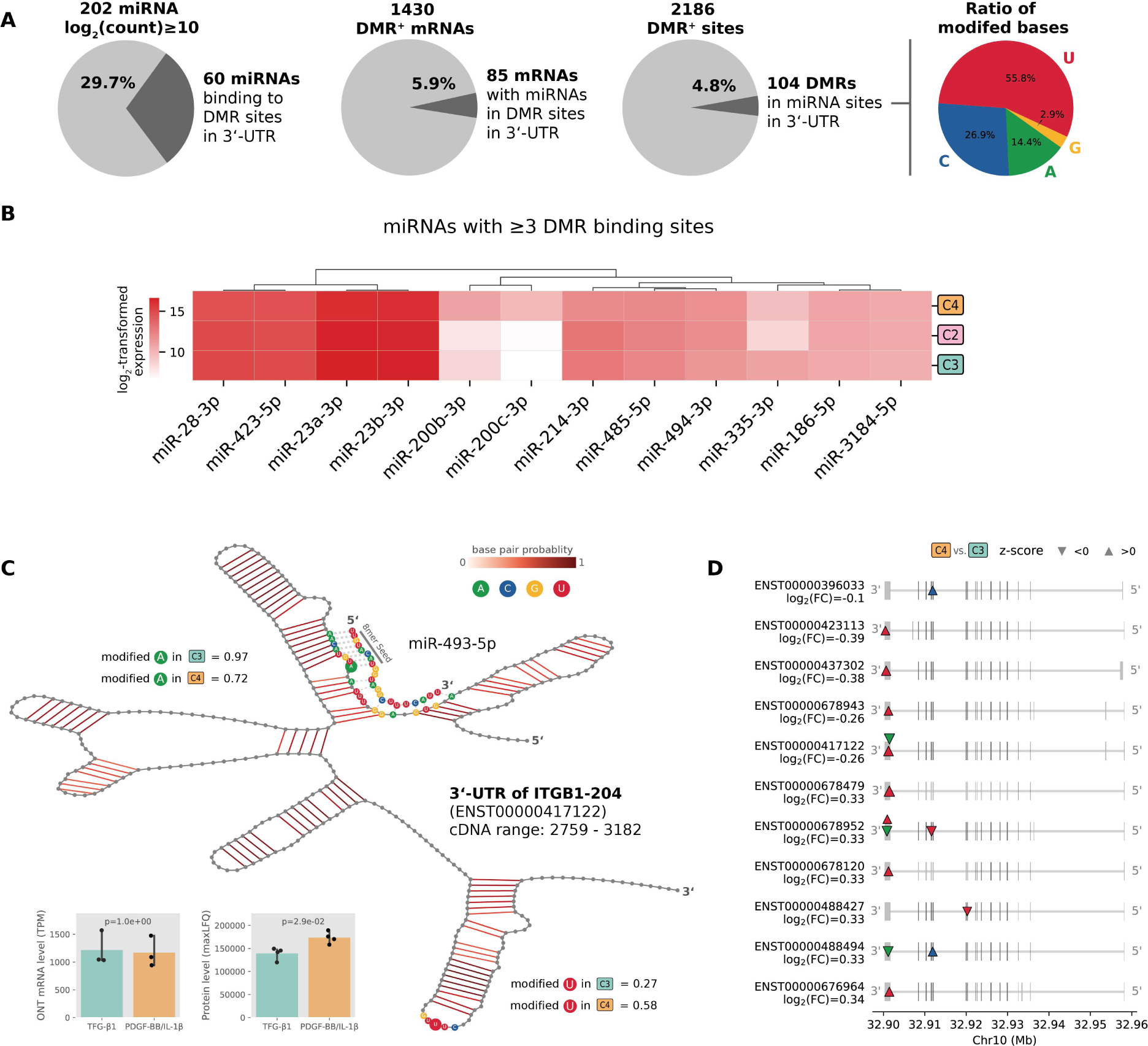
Potential binding of miRNAs at differentially modified RNA bases (DMRs). (**A**) Pie charts summarize highly expressed miRNAs (log_2_ count ≥ 10) predicted to bind DMRs via 8-mer seed sequences, the fraction of DMR-positive transcripts and DMRs within 3′ UTR miRNA binding sites, and the proportion of modified bases at these sites. (**B**) Expression pattern of highly expressed miRNAs with ore than three predicted DMR binding sites across three conditions (C2=TGF-β1 > PDGF-BB / IL-1β, C3=TGF-β1, C4=PDGF-BB / IL-1β). (**C**) *ITGB1* isoform ITGB1-204 illustrates a modified adenosine within a validated miR-493-5p binding site^38^ showing reduced modification in PDGF-BB / IL-1β–treated cells accompanied by increased ITGB1 protein levels. A second DMR was identified downstream, with enhanced uridine modification in C4. (**D**) Overview of DMRs across ITGB1 isoforms, with log_2_ fold change (FC; C4 vs C3) shown below each Ensembl transcript. Triangles indicate increased or decreased modification.

The highest number of DMR binding sites detected for miR-494-3p (n = 6), followed by miR-186-5p, miR-200b/c-3p, and miR-28-3p (n = 4) (**Figure 7B**). DMR-miRNA pairs were found for 85 mRNAs and 104 DMRs, thereby affecting about 6% of all DMR-positive transcripts. Of those, 58 sites have uridine modified, 28 cytidine, 15 adenosine, and 3 guanosine (**Figure 7A**). However, miRNA binding for only six genes had experimental evidence and a respective publication recorded in the miRTarBase database. In addition, the specific binding site of miR-493-5p at the unique DMR-site AA-A-UG of *ITGB1-204* in 3’-UTR (cDNA position 2859) was the only one experimentally confirmed^38^ but not yet recorded n miRTarBase **Figure 7C** illustrates a possible miRNA binding model postulating that miR-493-5p directly binds to a modified adenosine located in the AA-A-UG sequence. Another DMR site (GU-U-UC) is located about 230 nucleotides downstream at the tip of a hairpin loop. Interestingly, the mRNA levels similar in atheroprotective (TGF-1β; C3) and pro-atherogenic VMCs (PDGF-BB / IL1-β1; C4), yet integrin β1 protein levels appear to be higher in the latter VMC phenotype. In this model, elevated uridine modification in pro-atherogenic VMCs, particularly within a hairpin loop, could promote mRNA stability or reinforce RNA structural integrity. Conversely, decreased adenosine modification may reduce miRNA-binding efficiency, thereby limiting miRNA-mediated translational repression. Notably, the miRNA context-binding sequence is conserved across all *ITGB1* isoforms, where modified uridines or adenosines lie nearby (**Figure 7D**).

In order to investigate whether increased levels of miRNAs can influence RNA base modification, *e.g.* by decorating the modifiable site, we profiled the mirNOME in phenotypic transition conditions C2 to C5 (**Suppl. Table S19** and **S20**). In total, 131 miRNAs were significantly differentially expressed all four conditions, with the top four being miR-145-5p, miR-204-5p, miR-146a-5p, and miR-5585-3p. Thereby, miR-145-5p showed the greatest expression difference, exhibiting a ∼4-fold higher abundance in TGF-β1–treated VSMCs (C3) compared with PDGF-BB / IL-1β–treated VSMCs (C4) (**Suppl. Table S20**). This is consistent with prior studies demonstrating time- and dose-dependent downregulation of miR-145-5p under PDGF stimulation.^39^ However, neither for miR-145-5p nor the other differentially expressed mRNAs did we find evidence that miRNA binding directly affects RNA base modifications at specific miRNA binding sites.

## Discussion

Using Nanopore direct RNA-Seq, we performed a transcriptome-wide, untargeted screening of RNA base modifications in vascular smooth muscle cells (VSMCs) under atheroprotective (TGF-β1) and pro-atherogenic (PDGF-BB / IL-1β) treatment. Our main findings are that 1) up to 5% of mRNA bases appear to be modified (*e.g.* methylated). 2) Differences in mRNA base modifications between atheroprotective and atherogenic VSMC phenotypes often occur near stop codons and at the beginning of 3’-UTRs, specifically affecting uridines in accessible 3‘-UTR regions. 3) The G-Y-U-YY (with Y=U/C) is the most prominent motif found in ∼26% of isoforms. 4) Differentially modified RNA bases might be explained by global changes in RNA-modifying enzyme levels. 5) DMRs might be linked to translation activity, *e.g.* by regulating Poly(A) tail dynamics and miRNA binding.

These findings expand the understanding of VSMC phenotype switching in the context of atherosclerosis beyond transcriptional and epigenetic regulation.^5^ Although we were not able to detect traces of mRNA oxidation in oxidatively stressed VSMCs using Nanopore direct RNA-Seq, our multi-level analysis suggests that mRNAs can harbor multiple types of base modifications. Further, pro-inflammatory cytokines can influence the epitranscriptomic landscape by altering the expression levels of RNA-modifying enzymes. For example, nominally significant shifts in mRNA abundance for the pseudouridine writer *PUS7* and several ^5^C modifiers (*TET1 TET2 NOP2 NSUN2*) may explain the higher rate of uridine and cytidine modification in pro-atherogenic VMCs detected by xPore However, the molecular mechanisms and temporal dynamics underlying cytokine-induced regulatory programs that involve several classes of mRNA-modifying enzymes are largely unresolved. In particular, the signaling pathways and transcription factors that control the expression of specific mRNA-modifying enzymes, the stability of these enzymes under defined conditions, and the kinetics of base modification on endogenous mRNAs remain to be elucidated.

The most strongly enriched sequence context was the GY-U-YY motif (Y = pyrimidine base, C or U). To our knowledge, this motif has not been described so far. Our epitranscriptome-wide analyses suggest that modification of uridine may foster the PI3K/AKT signaling pathway triggered by PDGF stimulation^40^ by stabilizing transcripts essential for this pathway while destabilizing transcripts associated with antagonistic pathways. For example, many uridine-modified transcripts downregulated in pro-atherogenic VSMCs are under the control of the transcription factor TCF21, which normally suppresses PI3K/AKT signaling.^4^ TCF21 is known as a key driver of VSMC state transitions in atherosclerosis^2^ and the fact that many base-modified transcripts are regulated by TCF21 highlights the importance of mRNA base modifications in atherosclerosis.

To elucidate regulatory mechanisms underlying differentially modified RNA bases (DMRs), we investigated multiple possible consequences of DMRs on mRNA processing. We analyzed the impact of DMRs on mRNA splicing but did not find evidence for pathway-relevant differences in isoform usage in a DMR-dependent manner n fact, DMRs were often found in proximity to stop codons in agreement with previous studies^8,42^ indicating rather regulatory activity that might impact mRNA usage, mRNA folding, mRNA stability, or miRNA binding, and thus translational output.

First, we observed a negative correlation of protein abundance and poly(A) tail lengths, particularly for transcripts harboring DMRs, which is in line with previous study that also applied Nanopore direct RNA-Seq^42^ The potential contribution of DMRs highlights, indeed, the regulatory potential of these modifications.

Second, uridine modifications were enriched in predicted single-stranded RNA structure elements in 3’-UTRs, supporting the regulatory nature of these mRNA modifications, as these regions are potentially accessible for RNA-modifying enzymes and allow binding of proteins that exert post-transcriptional regulatory effects. The vast majority of significantly differential modifications affected uridine, likely representing pseudouridine (Ψ), which is the most common RNA modification. For Ψ, both stabilizing and destabilizing effects on mRNA were described^43^ and the examples of *NAMPT* and *FN1* hint that uridine modification in 3’-UTRs might be associated with enhanced mRNA stability and increased translational output (*cf NAMPT*), whereas uridine modification within coding sequence could potentially have opposite consequences (*cf. FN1*).

Third, it has been demonstrated that mRNA modifications can modulate miRNA binding when located in seed match regions in 3′-UTR miRNA binding sites.^44^ Notably, 60 out of 202 highly expressed miRNAs could directly bind to DMRs with their seed sequence, and modified uridine accounts for ∼56% of all affected DMR-sites. Although none of the predicted uridine modification-related miRNA binding sites were experimentally validated according to miRTarBase, hypothesize that of the identified uridine modifications in 3′-UTR can weaken miRNA binding and thereby prolong mRNA half-life and promote translational output. Nonetheless, additional in-depth studies are required to elucidate the regulatory implications of uridine modifications. Since some of the DMR-related miRNAs are directly related to SMC biology and atherosclerosis (*e.g.* miR-22, miR-143, miR-145, miR-200, miR-424)^45,46^ the notion of mRNA modifications affecting miRNA regulation might help to optimize miRNAs as novel therapeutics.

While this study is the first one that comprehensively combines investigations based on short-read sequencing, long-read direct RNA sequencing, as well as mass spectrometry, such multi-platform integration has technical limitations. For example, there is still a discrepancy between short-read RNA-Seq and long-read RNA-Seq with respect to determining expression estimates^47^ well mapping of reads. Further, quantification of long reads can be affected by incomplete transcript references, and recently, a large number of novel transcripts have been detected in a similar experimental setup to ours^19^ Despite the higher mapping accuracy of long-read sequencing, the limited throughput of MinION flow cells restricted ONT analyses to highly expressed genes that exhibit similar expression levels in two conditions, preventing differential modification testing for many transcripts. Consequently, insufficient coverage and limited replicates likely led to a considerable proportion of missed DMRs, which may explain the low number of detected modifications in oxidatively stressed VSMCs despite differential expression of multiple genes related to mRNA modification. Moreover, the time point of DMR assessment after cytokine stimulation might be crucial, as some RNA base modifications might lead to rapid mRNA degradation.

A further challenge in assessing the regulatory impact of DMRs on translational output is that the protein quantification relied on peptide-based measurements, which generally permit assignment at the gene level but not at the transcript level, thereby limiting the linkage of isoform-resolved DMRs to protein expression.

Another limitation is the use of the modification-type agnostic detection approach. Although k-mer context and enzyme-related gene expression suggest specific modification types (e.g., the DRACH motif for m6A), we did not detect strong signals of known sequence motifs at differential sites. Moreover, the ONT RNA002 chemistry precluded the se of ONT basecalling models for specific modifications, which are currently available only for RNA004 chemistry^48^ (*cf* https://genesilico.pl/modomics/tools). Consequently, accurate classification of individual, specific modification types will require dedicated technologies and experimental validation in future studies.

In conclusion, our modification-type-agnostic approach provides a comprehensive, transcriptome-wide view of the RNA modification landscape underlying VSMC phenotypic transitions. We show that distinct RNA modification types frequently co-localize within the same transcripts, particularly near stop codons, suggesting combinatorial regulatory effects processes such polyadenylation, mRNA stability, and interactions with regulatory proteins or miRNAs Moreover, our data indicate that mRNA base modifications especially uridine modifications act as post-transcriptional regulators that fine-tune cytokine-induced, transcription factor–specific signaling pathways and help maintain atherosclerosis-relevant SMC phenotypes. Finally, modification sites may serve as targets for miRNA-based therapeutic strategies, and base modifications may be detectable in circulating cell-free RNA, highlighting their potential as biomarkers of disease severity.

## Acknowledgments

Initial investigations of the ONT data and of xPore results were performed by Lalitha Kamada as part of her Master’s thesis, which was co-supervised by IW and Sven Rahmann and conducted within the Medical Systems Biology Group of Hauke Busch. We are further grateful to Jeanette Erdmann (deceased 2023), former head of the Institute on Cardiogenetics at the University of Lübeck, for supporting the research environment in which this work was conducted. This work used the high-performance computing infrastructure at the Research Center Borstel, Leibniz Lung Center.

## Supplemental Material

- Supplementary Methods (PDF)
- Supplementary Figures S1-S9 (PDF)
- Supplementary Tables S1-S20 (XLSX)

