## Supplementary Methods for "Epitranscriptomic profiling of VSMC phenotypes reveals uridine modifications linked to post-transcriptional regulation"

### *Cell culture of vascular smooth muscle cells*

In total, eight human primary vascular smooth muscle cells (VSMCs) were obtained from multiple commercial vendors (Thermo Fisher Scientific/Gibco, Cell Applications, Inc., and PromoCell), all of whom adhere to the principles of the Declaration of Helsinki (see Supplementary Table S1). VSMCs were cultured in M231 cell culture medium (Gibco M-213-500) with 1x Smooth Muscle Growth Supplement (SMGS) (Gibco S-007-25) at 37 °C and 5% carbon dioxide (CO2) in T75 or T175 flasks (Greiner). VSMCs were split at a maximum of 80% confluency until a maximum of eight passages. VSMCs were washed with 1x phosphate-buffered saline (PBS) before detachment with 1x trypsin/EDTA for up to 5 min at 37 °C. VSMCs were seeded on a collagen I hydrogel (Ibidi, 5 mg/mL, collagen type I, rat tail) for CellROX assays, Nanopore RNA sequencing, and Illumina RNA sequencing (see below). In brief, collagen I (Col I) was thawed at 4 °C at least two hours before use. A solution of 300 µL Col I hydrogel (1.5 mg/mL) was prepared by mixing the following components (stored at 4 °C) in the exact order on ice: 20 µL of 10x SMGS, 5 µL of 1 M NaOH (Roth K021.1), 81 µL dd$H_2O$ (Double-distilled, invitrogen 10977-035), 4 µL of $NaHCO_3$ (7.5%, w/v, Biochemka 71627), 100 µL M321 cell culture medium, and 90 µL of 5 mg/mL Col I. Pour slowly into a tube and mix carefully by pipetting up and down. The solution should have a pH of 6.5 - 7.5 (optimally 7.0). The freshly prepared Col I hydrogel was pipetted into each well/flask on ice. To enable polymerization, Col I hydrogel plates were stored at 37 °C for 30-60 min. In total, 0.2 mL and 1.8 mL of Col I hydrogel were used per well of a 48-well plate and a 6-well plate, respectively. After polymerization, $0.4\times10^5$ cells/well and $3.8\times10^5$ cells/well were seeded in a 48-well plate and a 6-well plate, respectively, and were incubated for 24 hours in M321 cell culture medium supplemented with 1% fetal bovine serum (FBS, BioSell FBS.GP.0500). In addition, 1 µM cytochalasin D (Sigma C2618) was added to inhibit VSMC contraction.[1] To induce the contractile phenotype of VSMCs, cells were incubated in fresh M321 medium supplemented with 1% FBS, 5 ng/mL human recombinant TGF-β1 (Transforming Growth Factor Beta 1, Peprotech 100-21C), and 1 µM cytochalasin D for four days. The transition from the contractile VSMC phenotype to a synthetic VSMC phenotype was achieved by replacing the old medium with fresh M321 medium (without cytochalasin D) supplemented with 1% FBS, 10 ng/mL human recombinant PDGF-BB (Platelet-Derived Growth Factor), and 10 ng/mL human recombinant IL-1β (Interleukin-1β). VSMCs were cultured for two days under these pro-inflammatory conditions.

### *Oxidative stress assays in vascular smooth muscle cells*

Two male human aortic VSMC lines and four technical replicates per VSMC line were used for oxidative stress assays: HAOSMC IV (Thermo Fisher Scientific, C-007-5C, LOT: 2164581) and HAOSMC 1596 (Cell Applications, Inc., 354-05a, LOT: 354-05a). Four (HAOSMC IV) or five (HAOSMC 1596) technical replicates were used for oxidative stress assays (cf. **Supplementary Table S1**). Each replicate value is the mean of four to five wells per technical replicate. VSMCs were seeded on a Col I hydrogel and incubated in M231 cell culture medium supplemented as described above in detail: 1) one day in M231 + 1% FBS, 2) four days with or without 5 ng/mL TGF-β1, and 3) two days with or without 10 ng/mL PDGF-BB and 10 ng/mL IL-1β. To induce measurable endogenous production of ROS, VSMCs were treated with 200 ng/mL PDGF-BB for 1 – 2 hours (referred to as PDGF boost), 30 µM CellROX® Green reagent (Invitrogen™, C10444), and 1 µg/mL Hoechst 33342 (Merck #14533-100MG) in M231 + 1% FBS. To further boost production of reactive oxygen species (ROS), VSMCs were exposed to ultraviolet (UV) radiation for 5 minutes. VSMCs without PDGF boost but with CellROX® Green reagent served as a negative control. N-Acetylcysteine (NAC), a potent antioxidant[2], served as the second control and was added two hours prior to and during the CellRox assay at a total concentration of 1 mM to quench ROS production. CellROX® Green fluorescent levels were measured with an immunofluorescence microscope (Keyence BZ-X800) using 10 µm z-stacks at 10-fold magnification and quantified with an in-house Python script. CellROX® Green fluorescent levels were normalized against Hoechst 33342 staining of nuclei.

***Quantification of the mitochondrial membrane potential***

The VSMC line HAOSMC IV was used to quantify the mitochondrial membrane potential (MMP) and the number of mitochondria (numMT) under different stimuli. Approximately $0.4\times10^5$ cells/well were seeded in 96-well plates (for MMP) or 8-well Lab Tek chamber slides (for numMT) previously coated with 1% porcine gelatine. After one day of incubation in M231 cell culture medium with SMGS, the medium was replaced with fresh M231 cell culture medium with 1% FBS. Cells were stimulated like the CellROX® assays as described above. To measure the MMP, cells were incubated with 200 nM TMRE (tetramethylrhodamine ethyl ester perchlorate, Invitrogen™ T669) in M231 + 1% FBS for 30 minutes at 37 °C and 5% CO2, washed with PBS, incubated with 1 µg/mL Hoechst 33342 for 20 minutes, washed twice with PBS, and fixated with cold 4% paraformaldehyde (PFA) for 20 minutes. To quench the MMP completely, 10 µM FCCP (Carbonyl cyanide-p-trifluoromethoxyphenylhydrazone; Sigma), a proton uncoupling agent, was added as a negative control. To check for the number of mitochondria (numMT), stimulated cells (see above) were fixated with cold 4% PFA for 10 minutes and stored at -20 °C until usage. On the day of staining, frozen cells on chamber slides were directly fixated in acetone-methanol (stored at -20°C) for 10 minutes. Cells were permeabilized with 0.1% Triton X-100 (Sigma 23.472-9) + 1% bovine serum albumin (BSA) for 30 minutes at room temperature (RT). Cells were blocked with 3% BSA in PBS for 1 hour at RT. Finally, cells were stained with an anti-GRP75 antibody (abcam ab53098, 1:1000) and an anti-β-Actin antibody (Sigma A5228, 1:500). Primary antibodies were visualized with Alexa Fluor 488 (Thermo Fisher Scientific, A11011) or Alexa Fluor 568 antibodies 488 (Thermo Fisher Scientific, A10037; 1:1000 dilution). TRME, GRP75, and Hoechst 33342 were quantified as described above (see CellROX® assays) using an immunofluorescence microscope (Keyence BZ-X800) and a Python script.

***Seahorse assays for vascular smooth muscle cell respiration***

The VSMC line HAOSMC IV was used for Seahorse assays. Approximately $0.4\times10^5$ cells/well were seeded in 24-well cell culture plates (Agilent #100777-004), leaving 4 wells blank. Cells were stimulated like the CellROX® assays as described above. Incubation with 200 ng/mL PDGF-BB was extended to 4 hours. Untreated cells served as the control. Five wells per condition were used as technical replicates. Seahorse experiments were repeated five times. XF24 sensor cartridges (Agilent #100850-001) were incubated overnight in Agilent calibrant solution at 37 °C under an ambient atmosphere. Approximately 1 hour before measurement, the cell culture medium was replaced by Seahorse XF DMEM assay (Agilent #103681-100) supplemented with 1 mM Pyruvate (Gibco), 4 mM L-glutamate (Gibco), and 10 mM Glucose (Sigma). To measure changes in oxygen consumption rate (OCR) and extracellular acidification rate (ECAR) using a Seahorse XF24 analyzer, 1.4 µM oligomycin (Sigma), 1 µM FCCP (Sigma), and 1.5 µM antimycin A/1.5 µM rotenone were added sequentially in each well. Each OCR measurement consisted of 1 min of mixing, 2 min of waiting, and 4 min of continuous $O_2$ level measurement. Cells were stained with Hoechst 33342 (Merck #14533-100MG) and counted to normalize OCRs. All Seahorse parameters (*e.g.*, basal respiration or ATP production) were calculated according to the manufacturer's instructions.

### ***Short-read RNA sequencing***

Short-read sequencing (NovaSeq6000, 2x100 bp) using the TrueSeq stranded mRNA library preparation kit without polyA enrichment was performed for eight biological replicates (see **Suppl. Table S1**) and five conditions (oxidative-stressed VSMCs (C1), oxidative stress-prone VSMCs (C2), atheroprotective VSMCs (C3), proatherogenic VSMCs (C4), and untreated VSMCs (C5)). Stimulation of VSMCs was conducted similarly to the CellROX® assays. Approximately $3.8\times10^5$ cells/well were seeded in 6-well plates that were covered with a Col I hydrogel (1.8 mL with 1.5 mg/mL). At least two wells per condition were prepared, and VMCS were harvested by incubating with 0.1% (w/v) collagenase type 2 in PBS (Cellsystems LS004176) for 15 minutes at 37 °C. Detached cells were centrifuged at 300×g for 5 minutes. The cell pellets were frozen at -80 °C until further usage. Total RNA (5 µg per reaction) was isolated using the innuPREP RNA Mini Kit 2.0 (Analytik Jena) according to the manufacturer's instructions. Depletion of ribosomal RNA (rRNA) was achieved with magnetic beads using the RiboMinus™ Eukaryote System v2 (Invitrogen A15026) according to the manufacturer's instructions. To remove all residues of the hybridization buffer, the magnet beads were washed twice instead of once. The rRNA-depleted RNA was concentrated from 300 µl supernatant with RNA clean XP beads (Beckman Coulter A63987). Initially, 660 µL of XP beads were added to a 1.5 mL tube and placed on a magnetic rack to allow separation. After removing 120 µL supernatant, the beads were resuspended and incubated at 65°C for 2 minutes. The bead suspension was added to the 300 µL RNA sample and mixed gently using a pipette. The mixture was incubated at RT for 15 minutes on a HoLa shaker to facilitate binding. Following this, the tube was placed on a magnetic rack to separate the beads, and the supernatant was removed and discarded. The beads were washed with 70% freshly prepared ethanol by rotating the tube 180° twice, followed by removal of the supernatant. The RNA-loaded beads were air-dried for 2–5 minutes before being resuspended in 30 µL RNase/DNase-free $H_2O$ or RNase/DNase-free TE buffer (Invitrogen 12090015). To elute the RNA, the sample was incubated at 65°C for 1 minute, followed by 4 minutes at RT. Beads were separated using a magnetic rack, and the supernatant containing the purified RNA was transferred to a new tube. RNA concentration was determined with the Qubit RNA HS Assay (Invitrogen Q32852) according to the manufacturer's instructions. The concentration of rRNA-depleted RNA was in a range of 2.7 ng/µL to 5.3 ng/µL.

### ***Short-read RNA sequencing data analyses***

Five conditions (oxidative stressed VSMCs (C1), oxidative stress-prone VSMCs (C2), atheroprotective VSMCs (C3), proatherogenic VSMCs (C4), and untreated VSMCs (C5)) were sampled with eight different VSMC cell lines as biological replicates, generating overall 40 RNA sequencing (RNA-Seq) datasets. These were analyzed using a community workflow from the snakemake workflow catalog (https://github.com/snakemake-workflows/rna-seq-kallisto-sleuth) version 2.8.4. This entails expression quantification with kallisto[3] and testing for differential expression using sleuth[4], for which the cell line was considered as a batch variable in the model. For raw and trimmed read QC, FASTQC was used (https://www.bioinformatics.babraham.ac.uk/projects/fastqc), and a combined QC report including mapping statistics was generated using MULTIQC[5]. For transcript annotation, Ensemble release 113 was used. Enrichment was computed via the workflow and used SPIA[6] internally for Reactome[7] and KEGG[8] pathways and GOATOOLS[9] for determining enriched Gene Ontology terms. Computation of switches between different isoforms of a gene was also computed via the workflow using the Biocondutor library IsoformSwitchAnalyzeR[10]. Principal component analysis (PCA) was performed for the 1,000 most variable genes after selecting the most highly expressed transcript per gene and keeping only transcripts with a minimum normalized log2 expression of 6.

***Long-read direct RNA sequencing by Oxford Nanopore Technologies***

The VSMC line HAOSMC IV was subjected to direct RNA-Seq using Oxford Nanopore Technologies (ONT) (**Suppl. Table S1**). Four conditions (oxidative stressed VSMCs (C1), oxidative stress-prone VSMCs (C2), atheroprotective VSMCs (C3), proatherogenic VSMCs (C4)) were sequenced. For this, approximately $5\times10^5$ cells were seeded into one T25 flask covered with 2 mL of 1.8 mg/ml Col I hydrogel. Stimulation, sample preparation, and RNA isolation were performed as described above for short-read RNA-Seq. To trigger endogenous ROS production, VSMCs were incubated with 400 ng/mL PDGF-BB for 3 hours. Approximately $10\times10^6$ stimulated cells per condition were used as input for direct RNA-Seq. Library preparation and direct RNA-Seq were conducted according to the ONT protocol SQK-RNA002 (version DRS_9080_v2_revQ_14Aug2019). About 150 – 200 ng rRNA-depleted RNA in 10 µL was used as input. A GridION device with FLO-MIN106 flow cells with more than 1000 active pores was used for sequencing. Direct RNA-Seq runs were stopped after 18 - 22 hours.

***ONT direct RNA sequencing data analyses***

ONT direct RNA-Seq data was processed using our own snakemake workflow available at https://github.com/iwohlers/2025_ont_rnaseq_vsmc, with tools installed in a reproducible fashion via conda using environment files provided as part of the workflow. The workflow uses guppy version 6.3.2 for basecalling and pycoQC[11] for quality control. Expression quantification was performed after mapping to hg38 using minimap2[12] with option '-x splice' followed by nanocount[11] with mRNA transcript annotation from Ensembl version 113 (http://ftp.ensembl.org/pub/release-113/gtf/homo_sapiens/Homo_sapiens.GRCh38.113.gtf.gz ). Mapping to Ensembl version 13 transcript sequences (http://ftp.ensembl.org/pub/release-113/fasta/homo_sapiens/cdna/Homo_sapiens.GRCh38.cdna.all.fa.gz) was performed with minimap2 using option '-x map-ont'. Primary mappings were used with nanopolish for event preprocessing (mode 'eventalign') and poly(A) tail length estimation (mode 'polya', coverage>10). For differential modification detection, xPore[13] version 2.1 was used with the nanopolish eventalign output, *i.e.*, modification positions refer to transcript coordinates. K-mer transcript positions from xPore (zero-based) were converted to relative genomic positions using pyensembl (https://github.com/openvax/pyensembl) and were converted to relative UTR and coding sequence (CDS) positions using the GFF3 file from Ensembl version 113.

Normalization of nanocount counts and testing of differential expression were performed with DESeq2[14], and an additional variance stabilization using rlog transformation was performed before principal component analysis, both as previously described for nanocount expression data.[11] These downstream analyses and visualizations are available as part of the FAIRDOMHub under https://fairdomhub.org/projects/461.

***Transcriptome-wide prediction of RNA structure elements***

RNA secondary structures were predicted and plotted with Python 3.11 using the ViennaRNA package library (https://github.com/ViennaRNA/ViennaRNA and https://www.tbi.univie.ac.at/RNA/ViennaRNA/refman/index.html; version 2.6.2) and the Forgi package library (https://github.com/ViennaRNA/forgi). In brief, 150-base mRNA sequences containing the modified base/k-mer were extracted (Ensembl version 113, see above). The first context sequence starts at -140 bases upstream and ends +10 bases downstream of the modified base. The sequence was shifted iteratively by two bases until the start base was -10 bases upstream, yielding up to 65 sequences per transcript and kmer. For each sequence, the centroid structure and corresponding minimum free energy (MFE in kcal/mol) were predicted. The centroid structure refers to a representative secondary structure derived from a Boltzmann-weighted ensemble of possible RNA conformations. The dot-bracket notation of the centroid structure (minus 10 bases at 5' and 3' end) was used to calculate the similarity score between all predicted structures using the Needleman-Wunsch algorithm (match score = 1, mismatch penalty = -1, gap penalty = -2). To speed up the algorithm, the similarity score was set to zero if the difference of MFE between the centroid structures was more than 10 kcal/mol. The similarity score was set to one if the centroid structures were the same. Centroid structures were clustered using hierarchical clustering implemented in SciPy. Clusters with the lowest mean MFE, the smallest MFE deviation, and the highest number of centroid structures were considered the best representatives of the RNA secondary structure. The five centroid structures with the lowest MFE of this representative cluster were used for further analysis. The Forgi package was used to convert the dot-bracket notation into secondary structure elements, such as unpaired nucleotides at the 5' end (f), stems or stacked bases (s, regions of contiguous canonical Watson-Crick base-paired nucleotides), hairpin loops (h, single-stranded regions enclosed by a stem), multiloop segments (m, a single-stranded region between two stems), interior loops (i, bulged out regions), and unpaired nucleotides at the 3' end (t). The over-representation analysis of secondary structure elements in kmers was conducted using the hypergeometric distribution function or Fisher's exact test calculated with SciPy.stats in Python. The background was overlapping counts of RNA secondary structure elements across all sequences.

***Prediction of RNA structure elements with miRNA binding***

Wide-window prediction (>350 nt) of RNA secondary structures was conducted with RNAplfold (version 2.7.0) with winsize=150, span=100, and default ulength=31. Alignment of the miRNA to the target sequence was performed by applying first local alignment (Smith-Waterman method without wobble bases; match=1, mismatch_penalty=-2, gap_penalty=-2) of the miRNA seed sequence and the complete transcript sequence, followed by global alignment (Needleman-Wunsch method with allowed wobble bases; match=1, wobble=1, mismatch_penalty=-1.5, gap_penalty=-2) of the complete mature miRNA sequence and the target sequence, which consists of the miRNA binding region extended by a 20 base upstream window. The RNA secondary structure of ITGB1-204, together with miR-493-5p, was plotted as a force-directed graph (Kamada-Kawai layout) with the Python package networkx. Base pairs were created if the base pair probability was above 0.5, and pseudoknots were excluded. Hairpin structures were kept only if at least three consecutive stacked bases were present in the stem region.

***Small RNA sequencing***

For the identification of highly expressed miRNAs, small RNA sequencing was performed for two technical replicates of four conditions (oxidative stress-prone VSMCs (C2), atheroprotective VSMCs (C3), proatherogenic VSMCs (C4), and untreated VSMCs (C5)), respectively. Approximately $3.8 \times 10^5$ cells/well (HAOSMC IV) were seeded in 6-well plates that were covered with a Col I hydrogel (1.5 mg/mL). VSMCs were stimulated as described above for short-read RNA-Seq. Cells were harvested and kept in TRIzol™ reagent (500 µL/well, Invitrogen 15596-018). Total RNA was extracted using the RNeasy Plus kit (Qiagen 74134). Adaptors were ligated to the 3' and 5' ends of small RNAs, respectively. After hybridization with reverse transcription primer, the first-strand cDNA was synthesized. The double-stranded cDNA library was generated through PCR enrichment. After purification and size selection, libraries with insertions between 18~40 bp were obtained. These libraries were checked with Qubit and real-time PCR for quantification and a bioanalyzer for size distribution detection. Quantified libraries were pooled and sequenced on an Illumina sequencer, targeting 20 million single-end reads of length 50 bases.

***Small RNA sequencing data analyses***

Small RNA-Seq reads were trimmed using cutadapt[15] to identify and trim up to five adapters and remove sequences (i) not containing any adapter, (ii) containing any N's, (iii) shorter than 16 and longer than 28 bases (since no target sequences have this size) (parameters "-a AGATCGGAAGAGCACACGTCT -g GTTCAGAGTTCTACAGTCCGACGATC --max-n 0 --discard-untrimmed --minimum-length 16 --maximum-length 28 -n 5"). Trimmed reads were mapped using kallisto[4]. Towards this, mirBase[16] release 22.1 mature miRNA sequences were downloaded from https://www.mirbase.org/download/CURRENT/mature.fa and indexed with k-mer size 13. Subsequently, reads were mapped using kallisto quant with fragment length 20 (-l 20) and length standard deviation 5 (-s 5) as well as parameters "--single --single-overhang".

We investigated whether the modified base is part of miRNA seed regions, for which we considered 7mer-m8 sites, *i.e.*, bases 2-8 of the miRNA need to match the reverse complement of the region covering the modified base, which includes 8mer sites having an adenosine as the first base at the target site. The seed regions were obtained from Targetscan[17] (https://www.targetscan.org/vert_80/vert_80_data_download/miR_Family_Info.txt.zip).

Annotation of publication-supported miRNA-target interactions was obtained from MirTarBase[18] (https://mirtarbase.cuhk.edu.cn/~miRTarBase/miRTarBase_2025/cache/download/10.0/MicroRNA_Target_Sites.csv).

***Proteomics***

Quantification of proteins was performed for TGF-β1 (atheroprotective; C3) and PDGF-BB (proatherogenic; C4) stimulated VSMCs using mass spectrometry with four technical replicates each. Proteins (~20 – 44 µg per cell replicate) were extracted with TRIzol according to the manufacturer's recommendation. Protein concentration was determined via µBCA assay (Thermo Scientific 23235) with BSA as a standard. Equal amounts (4 µg) of individual samples were treated with benzonase (0.125 U/µg protein) and subjected to sample preparation, which includes reduction and alkylation, protein precipitation on SP3 beads, and digestion with Trypsin/LysC Mix (1:25) overnight at 37 °C. Resulting peptides were subjected to mass spectrometry measurement on a HPLC-ESI-MS/MS system (HPLC=high pressure liquid chromatography, Ultimate 3000 UPLC system; ESI=electrospray ionization) coupled to an Orbitrap Exploris™ 480 mass spectrometer. MS data were analyzed and normalized with the Spectronaut software (https://biognosys.com/software/spectronaut/) using the Uniprot database limited to human entries (v.2023_12). In total, 4,993 proteins were identified with a minimum of two peptides per protein. Only unique and razor peptides were considered. Protein levels represent maxLFQ values (label-free quantification by delayed normalization and maximal peptide ratio extraction). The statistical analysis was performed on the peptide level using the PECA[19] package (version: 1.40.0). A ROTS test (ROPECA approach - reproducibility-optimized peptide change averaging) implemented in the PECA package was used to perform the statistical analysis (https://doi.org/10.1371/journal.pcbi.1005562). Protein set enrichment analysis was conducted using the Enrichr[20] API, with all identified proteins serving as the background (proteins with ≥2 fragments).
