## Supplementary Figures S1-S9 for "Epitranscriptomic profiling of VSMC phenotypes reveals uridine modifications linked to post-transcriptional regulation"

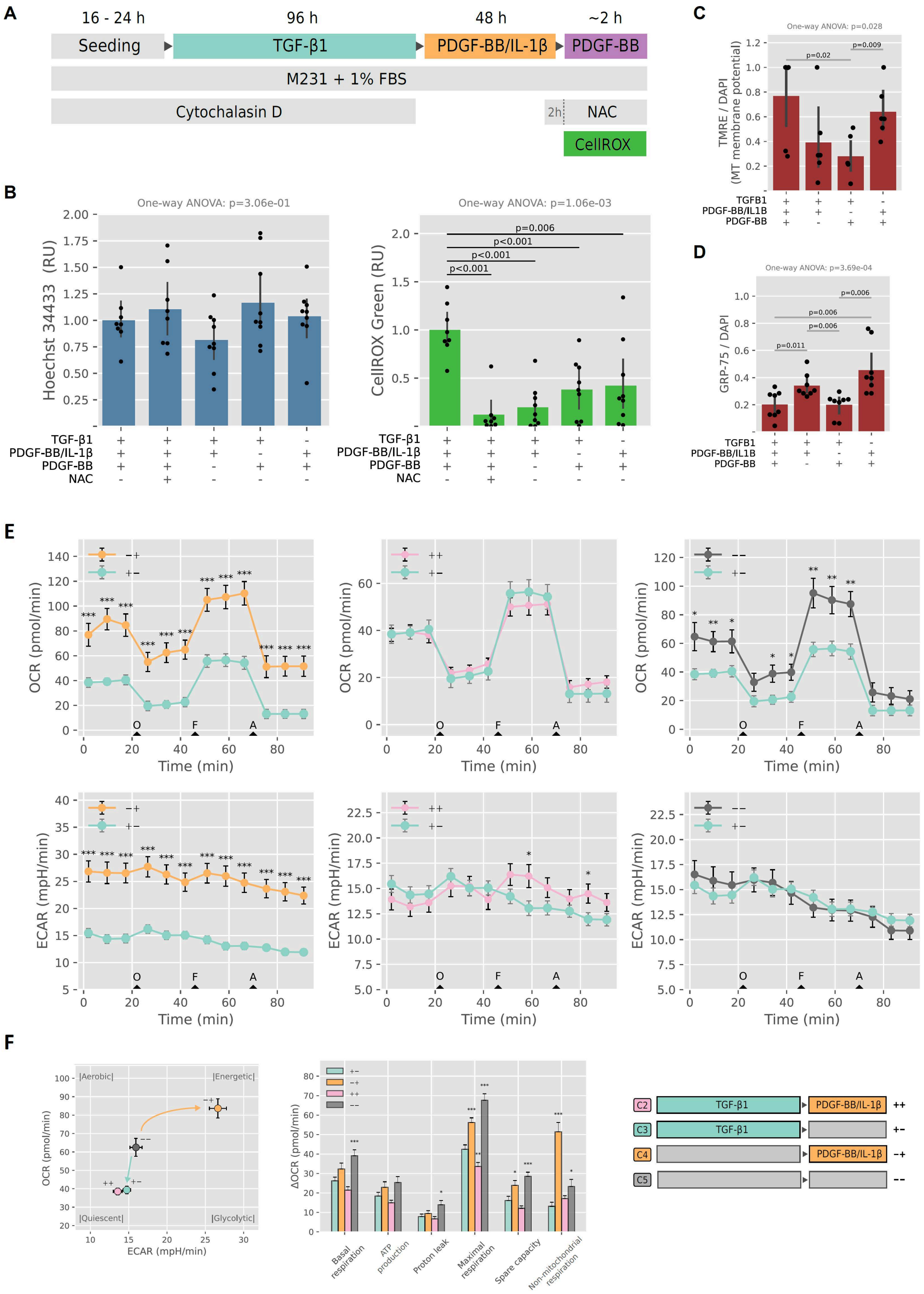


**Figure S1**: Oxidative stress and mitochondrial activity in vascular smooth muscle cells (SMCs) under atheroprotective and pro-atherogenic conditions. (**A**) VSMC culture and treatment scheme (FBS, fetal bovine serum; NAC, N-acetylcysteine). (**B**) Endogenous oxidative stress assessed by CellROX fluorescence, normalized to Hoechst 33342, following treatment with TGF-β1 (5 ng/mL), PDGF-BB (10 ng/mL) / IL-1β (10 ng/mL), or high-dose PDGF-BB (200 ng/mL). Two VSMC lines with n=4 (HAOSMC IV) or n=5 (HAOSMC 1596) technical replicates were used. NAC (1 mM) was used to reduce oxidative stress. (**C**) Mitochondrial (MT) membrane potential measured by TMRE fluorescence; (**D**) mitochondrial abundance assessed by GRP75 immunostaining. (**E**) Oxygen consumption rate (OCR) and extracellular acidification rate (ECAR) measured by Seahorse analysis in VSMCs treated with TGF-β1 (+−), PDGF-BB / IL-1β (−+), both growth factors (++), or untreated (−−). Approximately 1 hour before measurement, the cell culture medium was replaced by Seahorse XF DMEM assay

(Agilent #103681-100) supplemented with 1 mM Pyruvate (Gibco), 4 mM L-glutamate (Gibco), and 10 mM Glucose (Sigma). To measure changes in oxygen consumption rate (OCR) and extracellular acidification rate (ECAR) using a Seahorse XF24 analyzer, 1.4 µM oligomycin (O), 1 µM FCCP (F), and 1.5 µM antimycin A/1.5 µM rotenone (A) were added sequentially in each well. (**F**) Seahorse parameters were calculated according to the manufacturer's instructions. Each replicate value is the mean of three to five wells of Seahorse plates. In **C** - **F**, one VSMC line with n=6 (TMRE), n=7 (GRP-75), and n=5 (Seahorse) technical replicates was used (cf. **Suppl. Table S1**). In **B** - **F**, statistical analysis was performed by one-way ANOVA followed by unpaired Student's t-tests where applicable ($p < 0.05$). In **F**, statistical significance was calculated using a two-sided Student's t-test per time point.

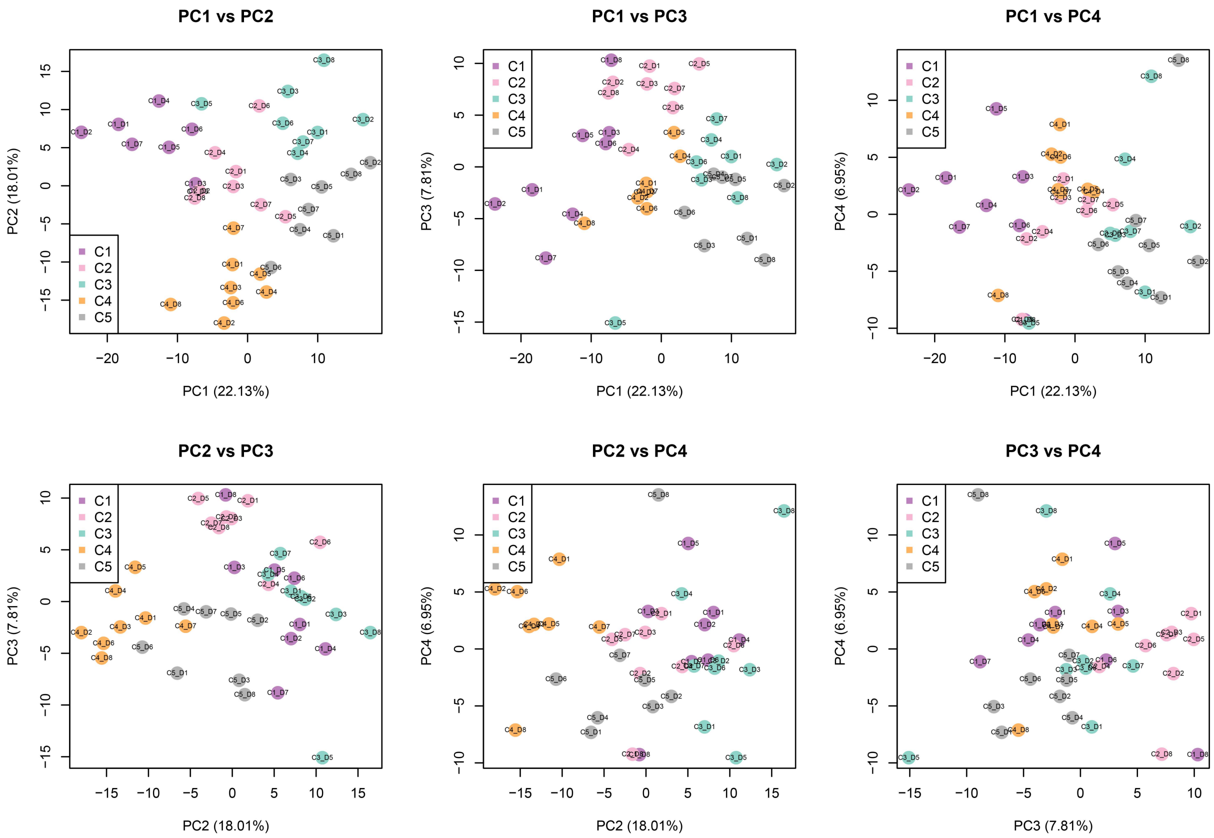


**Figure S2***:* Principal component-based transcriptome profiling of VSMC phenotypic conditions C1 to C5 based on Illumina short-read RNA sequencing of eight VSMC cell lines as biological replicates. Shown are all combinations of the principal components (PC) 1 to 4 for the 1,000 most variable transcripts after keeping the on average most highly expressed transcript per gene if its normalized and batch-corrected log2 expression is larger than or equal to 5 in all samples.

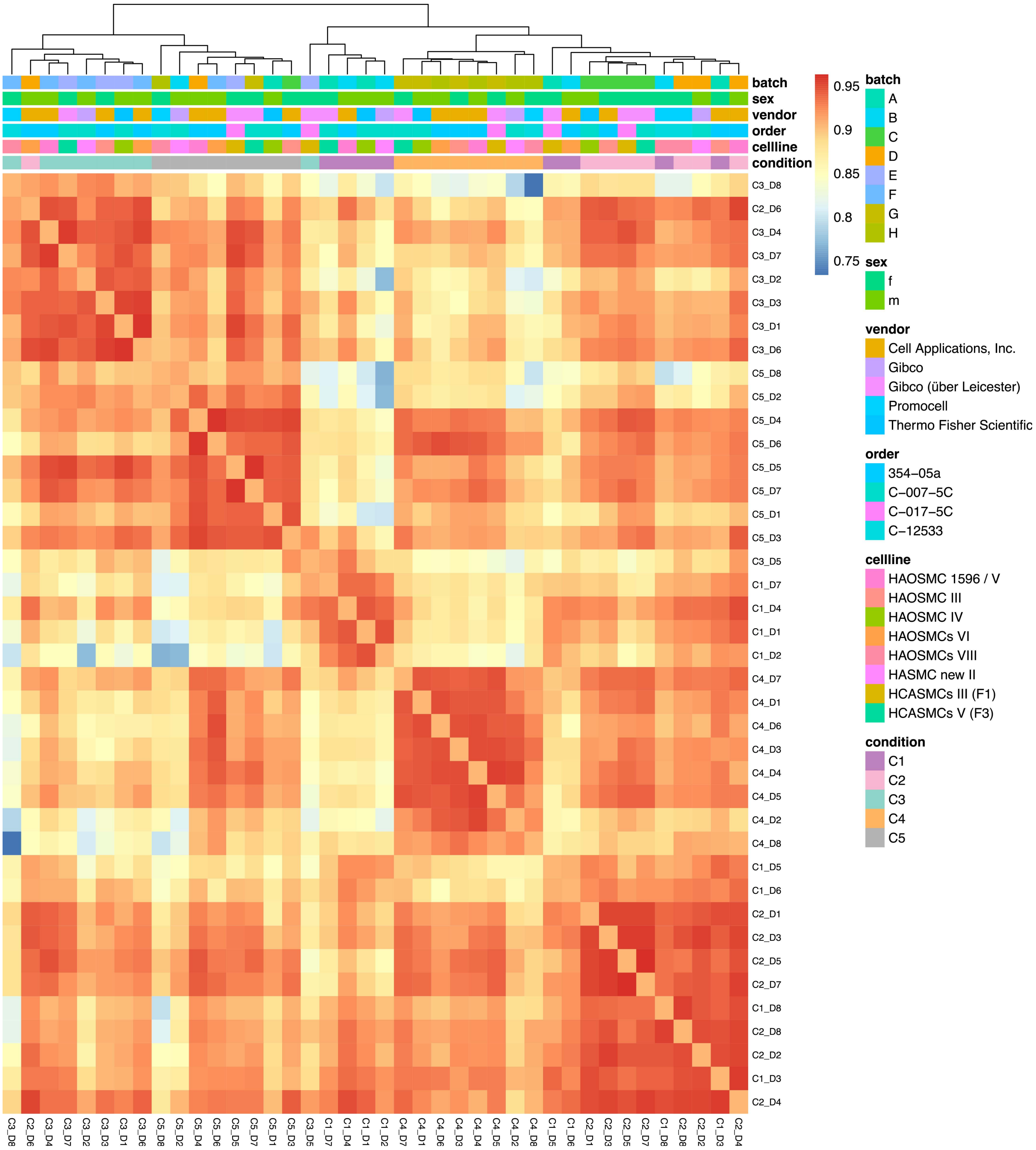


**Figure S3**: Sample-to-sample correlation heatmap based on pairwise Pearson correlations calculated from the 1,000 most variably expressed genes according to short read-based, log2-transformed expression after normalization across samples. The hierarchical clustering uses as distance 1 – correlation. Typically, expression profiles of samples from the same condition are most strongly correlated. Expression profiles of oxidative stress samples (C1) tend to be less correlated with the profiles of samples of other conditions.

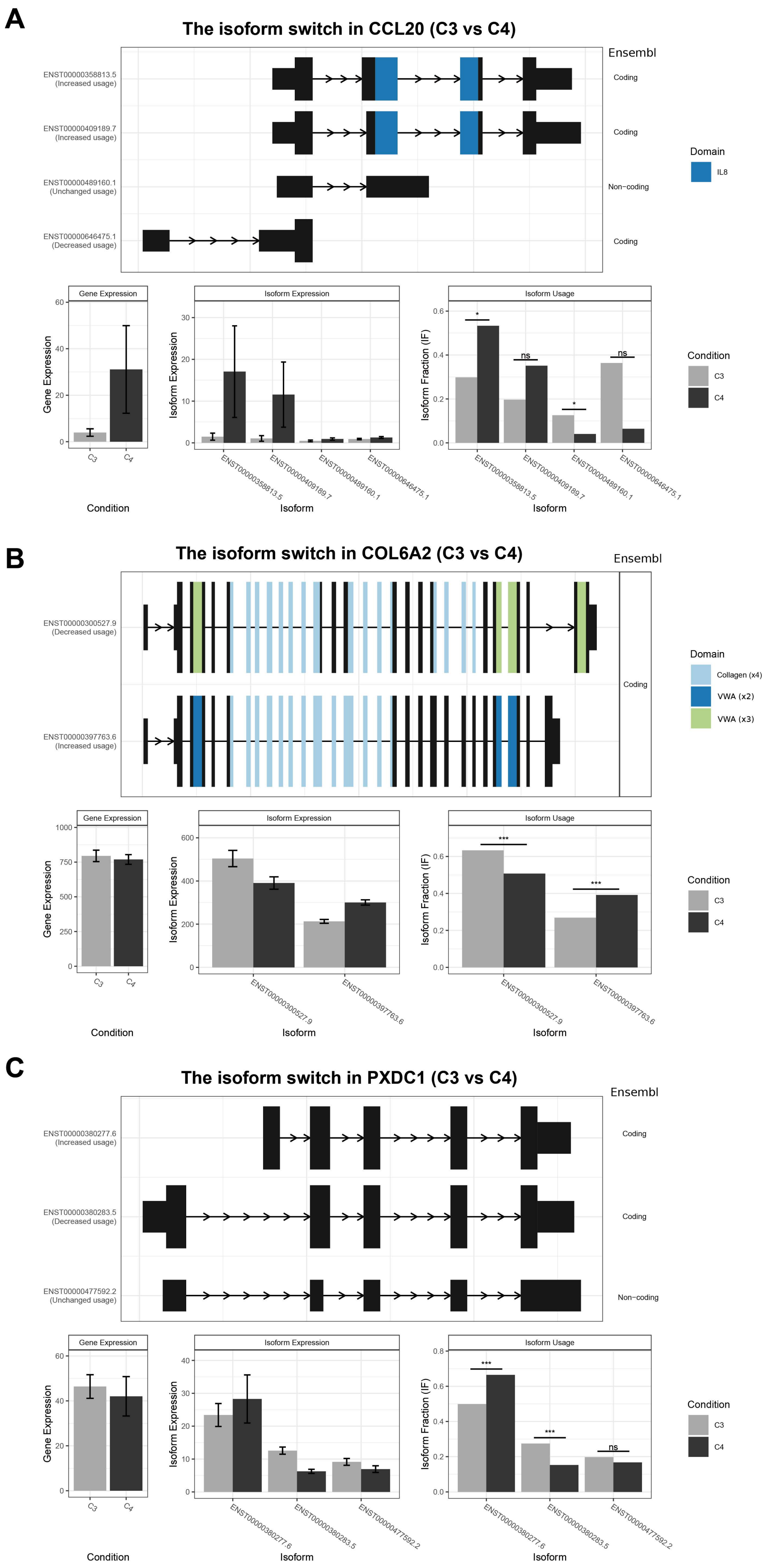
A
The isoform switch in CCL20 (C3 vs C4)
Ensembl
ENST00000358813.5 (Increased usage)
ENST00000409189.7 (Increased usage)
ENST00000489160.1 (Unchanged usage)
ENST00000646475.1 (Decreased usage)
Coding
Non−coding
Domain
IL8
Gene Expression
Isoform Expression
Isoform Usage
Isoform Fraction (IF)
Condition
Isoform
C3
C4
ns
B
The isoform switch in COL6A2 (C3 vs C4)
ENST00000300527.9 (Decreased usage)
ENST00000397763.6 (Increased usage)
Collagen (x4)
VWA (x2)
VWA (x3)
C
The isoform switch in PXDC1 (C3 vs C4)
ENST00000380277.6 (Increased usage)
ENST00000380283.5 (Decreased usage)
ENST00000477592.2 (Unchanged usage)

**Figure S4**: Isoform-usage analysis of (A) *CCL20*, (B) *COL6A2*, and (C) *PXDC1* from Illumina RNA-Seq data derived from vascular smooth muscle cells treated with atheroprotective growth factors (TGF-β1; C3) or pro-inflammatory factors (PDGF-BB / IL-1β; C4). Four of the overall five isoforms of *CCL20* are shown in A. Transcript ENST00000358813.5, ENST00000409189.7, and ENST00000646475.1 are annotated as protein-coding in Ensembl, although predicted non-coding from the coding potential assessment tool (CPAT). The first two include an IL8 domain, shown in blue. Both show a much higher expression than other isoforms in C4. Transcript ENST00000489160.1 is annotated as a retained intron. Significance is computed within the tool SwitchAnalyzeR.

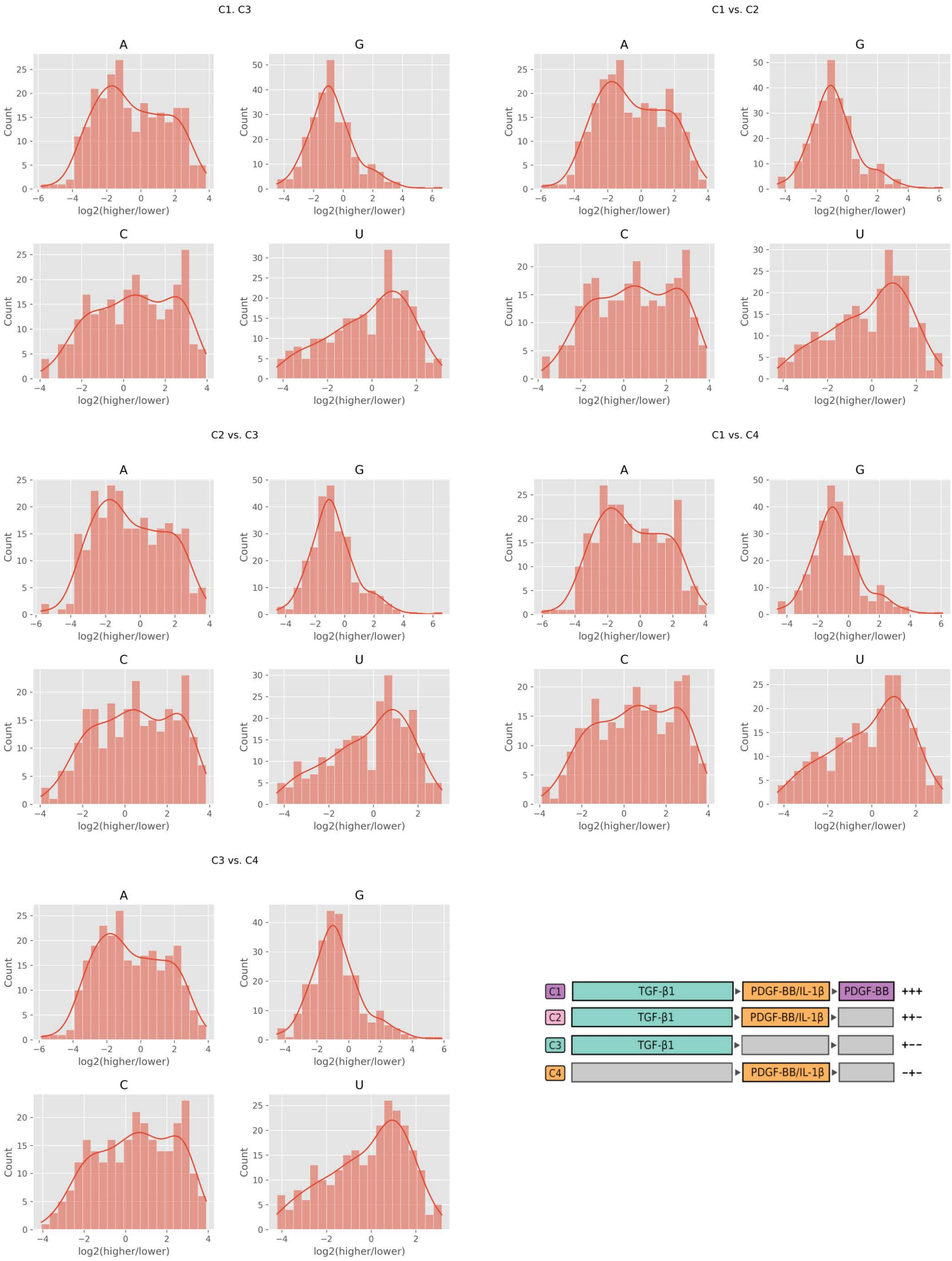


**Figure S5:** Histograms of xPore modification modes ("higher" vs. "lower") across condition C1 to C4 for each RNA base (A, G, C, and U) before filtering xPore data (*i.e.,* majority voting). The distribution pattern may reflect different base modifications. For example, modifications in G tend to result in lower raw Nanopore signals, while in A or C, at least two distinct base modifications can be suspected.

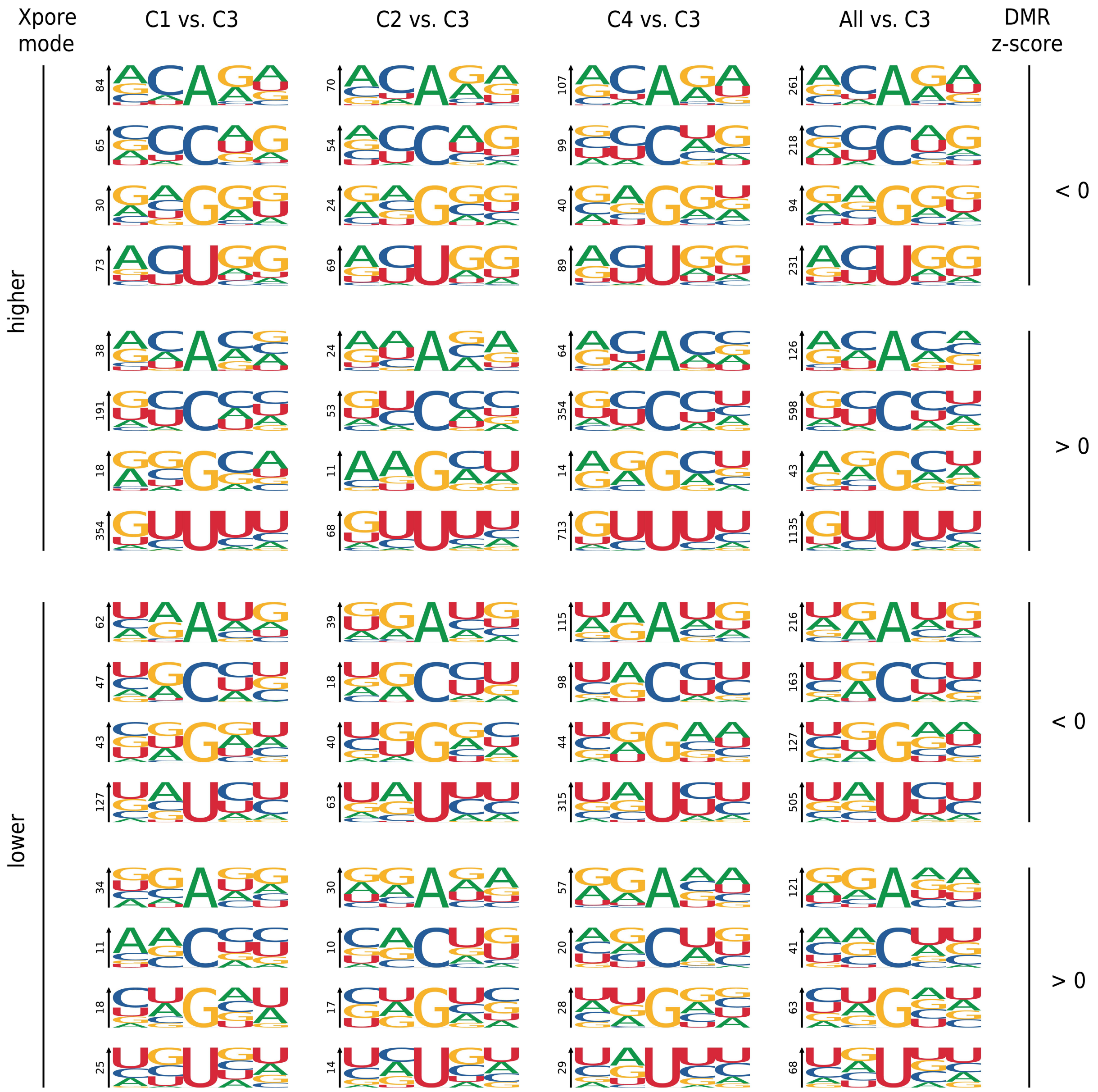


**Figure S6:** Number of significant sites as 5-mer motifs with differential modification rates more than 25% in comparisons with atheroprotective condition C3 (TGF-β1-treated smooth muscle cells). The sites are stratified by xPore mode ("higher" or "lower") and direction of modification change (reflected by z-score less than or greater than 0). Each 5-mer motif is generated across the sites with the respective mode and direction, with the number of significant sites denoted. Most prevalent are uracil modifications, and most prominent is the motif GUUUU across all comparisons (C1, C2, and C4; PDGF-BB / IL-1β-treated smooth muscle cells) with C3.

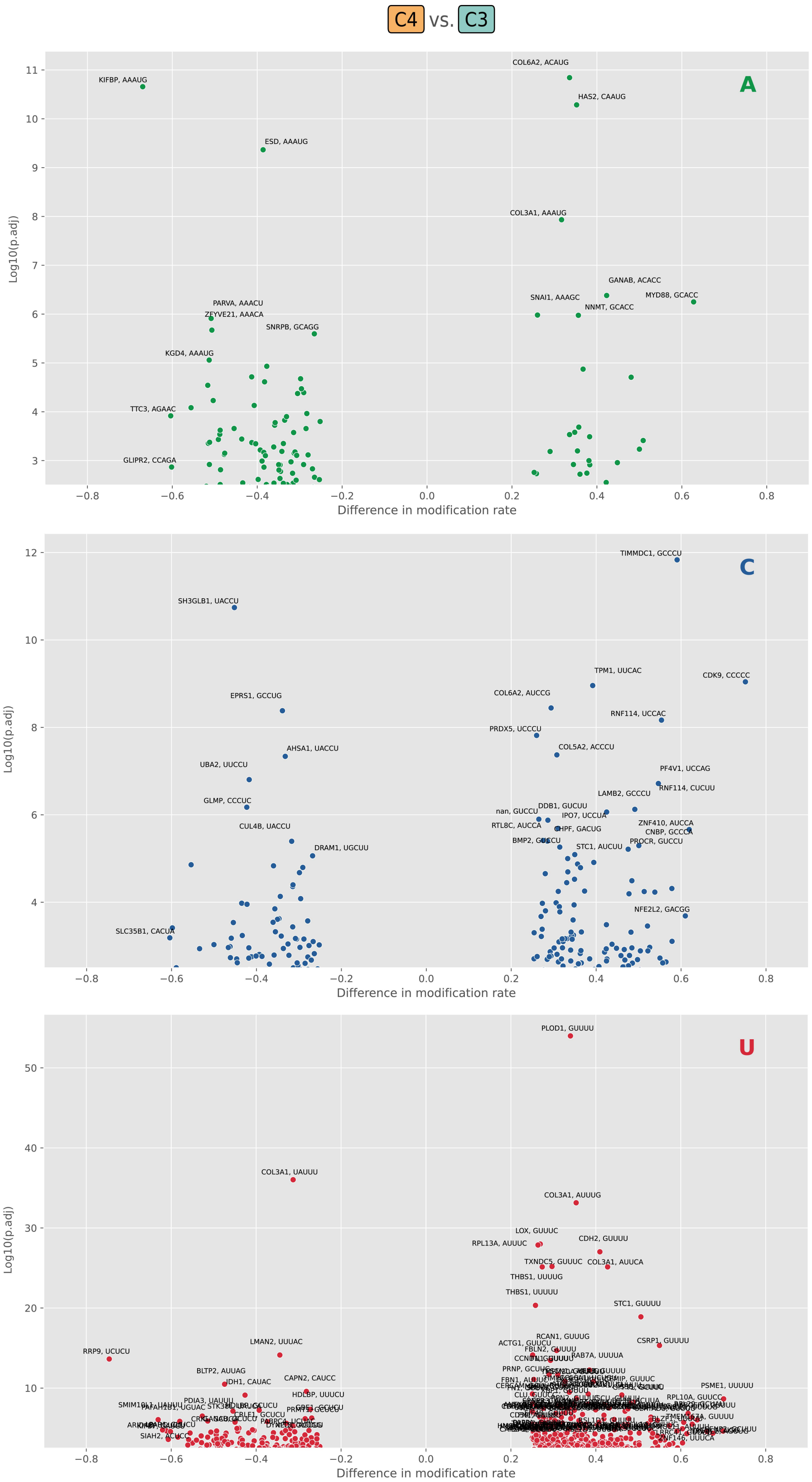


**Figure S7**: Volcano plots of differentially modified RNA bases in vascular smooth muscles treated with atheroprotective growth factors (TGF-β1; C3) or pro-inflammatory (PDGF-BB / IL-1β; C4) cytokines.

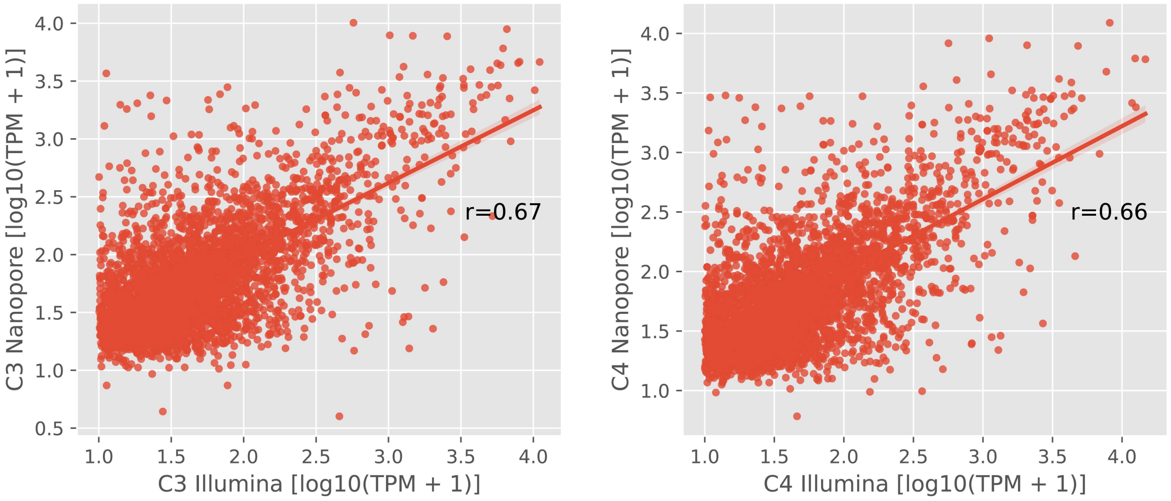


**Figure S8**: Correlation between Nanopore and Illumina log10-transformed mRNA levels with >10 TPM. Expression levels are means of three technical replicates (Nanopore) or levels derived from one Illumina sequencing run with the same SMC line used for Nanopore sequencing (HAOSMC IV; cf. Suppl. Table S1). The Pearson's correlation coefficient (r) is shown for VSMCs treated with atheroprotective TGF-β1 (left panel; C3; 3,843 transcripts) or pro-inflammatory PDGF-BB / IL-1β (right panel; C4; 3,808 transcripts). The p-value was <0.001 for both conditions and calculated with a two-sided t-test as implemented in the SciPy stats package.

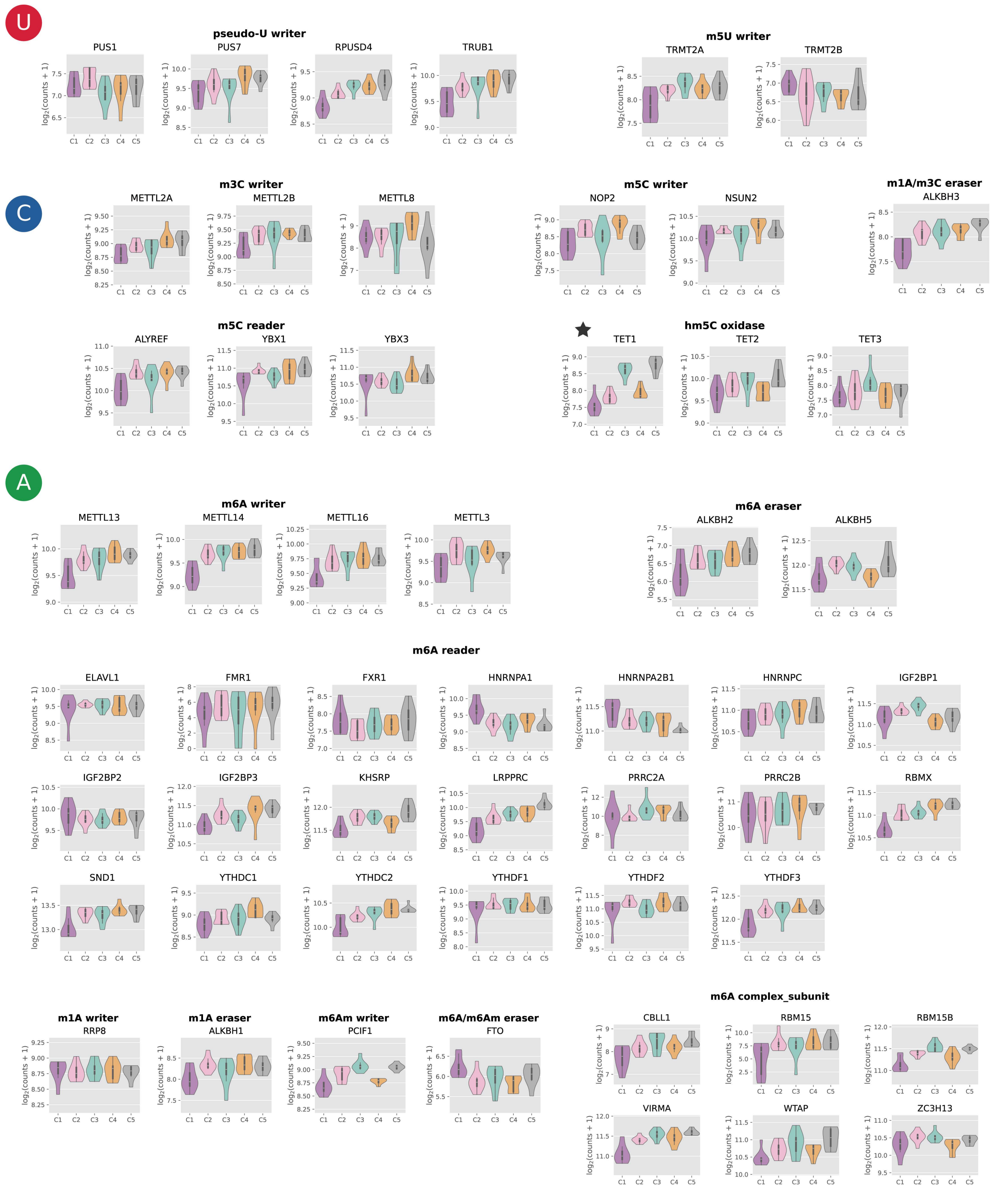


**Figure S9:** Gene expression of genes encoding proteins that alter RNA base modifications under conditions C1 to C5. A detailed description of conditions can be found in the main *Figure 1*. Asterisks indicate genes that are significantly differentially expressed in the comparison C3-C4 (p.adj. < 0.05).
